# A Sensitivity Analysis of Methodological Variables Associated with Microbiome Measurements

**DOI:** 10.1101/2023.12.12.571292

**Authors:** Samuel P. Forry, Stephanie L. Servetas, Jennifer N. Dootz, Monique E. Hunter, Jason G. Kralj, James J. Filliben, Scott A. Jackson

## Abstract

The experimental methods employed during metagenomic sequencing analyses of microbiome samples significantly impact the resulting data and typically vary substantially between laboratories. In this study, a full factorial experimental design was used to compare the effects of a select set of methodological choices (sample, operator, lot, extraction kit, variable region, reference database) on the analysis of biologically diverse stool samples. For each parameter investigated, a main effect was calculated that allowed direct comparison both between methodological choices (bias effects) and between samples (real biological differences). Overall, methodological bias was found to be similar in magnitude to real biological differences, while also exhibiting significant variations between individual taxa, even between closely related genera. The quantified method biases were then used to computationally improve the comparability of datasets collected under substantially different protocols. This investigation demonstrates a framework for quantitatively assessing methodological choices that could be routinely performed by individual laboratories to better understand their metagenomic sequencing workflows and to improve the scope of the datasets they produce.

## Introduction

The democratization of access to next-generation sequencing (NGS) has made this technology the workhorse of modern microbiology. NGS-based metagenomic sequencing (MGS) enables analysis of the genomic content of microbial communities, reporting both the taxonomic identification and count of sequencing reads and even enumeration of constituent microbes. In recent years, MGS has increasingly found its way into the clinic and other regulated spaces such as quality control testing where the measurement results can have important consequences.(1–3) Despite widespread adoption of MGS capabilities, the analytical performance remains uncertain, particularly when comparing between protocols.(4)

While MGS measurements may appear straightforward, the results actually depend on a complex series of steps and sample manipulations, typically including: sample storage and transport, DNA extraction and purification, DNA fragmentation and barcoding, library preparation and NGS, bioinformatic analysis to identify and enumerate unique microbial signatures, and taxonomic assignment against a database, as well as additional bioinformatic analyses dependent on particular experimental designs and project goals. Further, within each of these steps, many methodological choices must be specified for the particular protocol employed. Each of these steps, as well as the individual methodological choices, can introduce bias and variability that then accumulate through the measurement process. (5–7) The methodological contributions of bias and variability can be visually depicted using an Ishikawa diagram (Figure 1).

**Figure 1:**
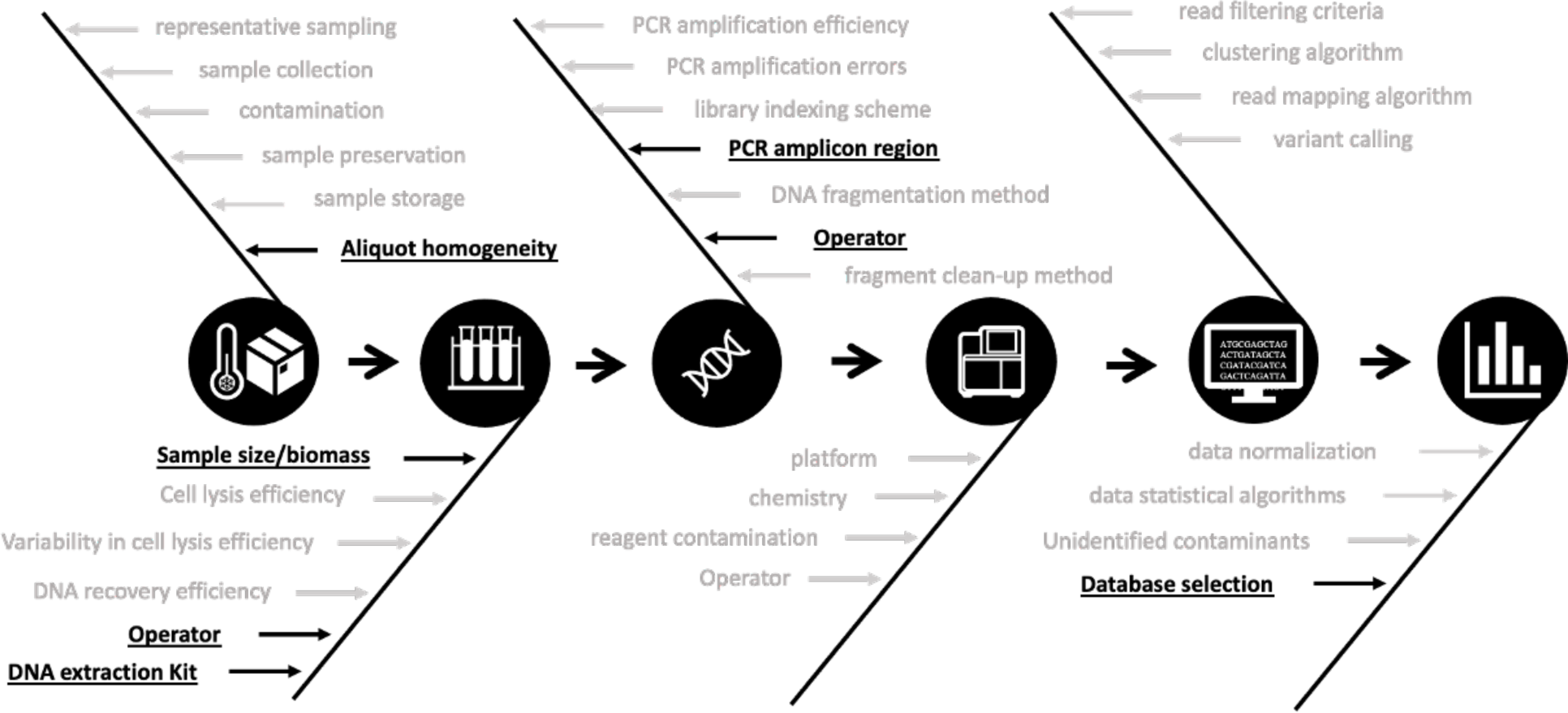
Ishikawa diagram of bias associated with an amplicon metagenomic measurement workflow. The central image depicts a typical metagenomic analysis: sample handling and storage, DNA extraction, sequencing library prep, next-generation sequencing, sequence data processing, and bioinformatic analysis. At each step, some of the various methodological choices that can introduce variability and bias are listed. Bold and underlined steps indicate methodological choices explicitly varied in the current study.

In general, MGS datasets exhibit high precision (low variability) for locked-down workflows (8, 9), and this makes MGS particularly useful within a research project where protocols can be easily controlled. Given the myriad options for any single step (as well as a dynamic field where procedures continue to evolve quickly), it is unlikely that two different labs would implement the exact same measurement workflow from start to finish without substantial coordination. Hence, the lack of reproducibility between laboratories is perhaps not surprising even when the results are highly reproducible within a laboratory.(10) While there are limited opportunities through the MGS workflow for intermediate characterization of the impact of each methodological decision, there are experimental designs aimed at assessing the effect sizes of particular methodological decisions at the conclusion of the MGS measurements.(11) One commonly used approach is a full factorial design, evaluating at least two specified options (‘Level’) for each tested methodological parameter.(12–14) This experimental design allows for a rigorous statistical assessment of the effect of individual parameters and provides quantitative data for direct comparisons.

We previously described the Mosaic Standards Challenge (MSC), an international interlaboratory study designed to assess the impact of methodological variability on MGS results.(15) The MSC employed five biologically distinct human fecal reference materials and a comprehensive standardized metadata reporting sheet that allowed participants to share exhaustive details about the protocols and methods used for analyzing the fecal materials. The MSC employed an “open protocol” design that allowed (and even encouraged) participants to follow divergent protocols as defined for their lab’s routine MGS measurements. This strategy showcased the diversity of methodologies commonly employed across MGS measurement labs and highlighted the challenge of comparing MGS results between different protocols. In the current effort, we used the same five fecal samples from the MSC, but employed a fully specified experimental design where a small subset of methodological choices were systematically varied while the rest of the MGS measurement workflow was held constant.

The goal of this work was to demonstrate a rigorous experimental approach for evaluating bias in metagenomics workflows. Specifically, we used a full-factorial design (plus replication) for each fecal sample (n=5) to quantify the effects of different operators (n=2), reference material lot (n=2), DNA extraction kit (n=2), marker gene target (n=2), and reference database (n=3). The magnitude of effect for each selected methodological choice was quantified, and these main effects were compared between protocol choices and to the biological variability between fecal samples. Ratiometric analysis was employed to account for MGS measurement compositionality. In combination with spike-in species added to each fecal sample, this analysis allowed direct comparisons between samples. Among the methodologies explored, the largest effects were observed for extraction kit selection, where the effect varied substantially between individual taxa, even within taxonomic clades.

## Results and Discussion

### Metagenomic Sequencing Methodologies and Experimental Design

Using an Ishikawa diagram, a subset of parameters were selected (shown in bold and underlined) for this initial proof-of-concept study (Figure 1). The experimental design included parameters that were denoted ‘biological’ (i.e., actual biological differences between disparate stool samples) or ‘methodological’ (i.e., bias associated with the measurement protocol) to allow for direct comparison of the variability from biological differences to the variability due to methodological differences. For this study, 5 unique human fecal samples were selected representing the biological variability. For each sample, the methodological parameters of operator (n = 2), lot (n = 2), extraction kit (n = 2), 16S variable region (n = 2), database (n = 3), were chosen (with the indicated number of levels). Thus, 240 MGS datasets (270 with replicates) were generated from 48 distinct workflows (Figure 2). The orange and blue arrows in Figure 2 at the top and bottom arms of the study, respectively, represented orthogonal workflows where unique methods were chosen for each parameter. The two orthogonal workflows were replicated to assess precision, and the orange and blue lines were used here and in subsequent figures to showcase data originating from the two completely distinct workflows.

**Figure 2:**
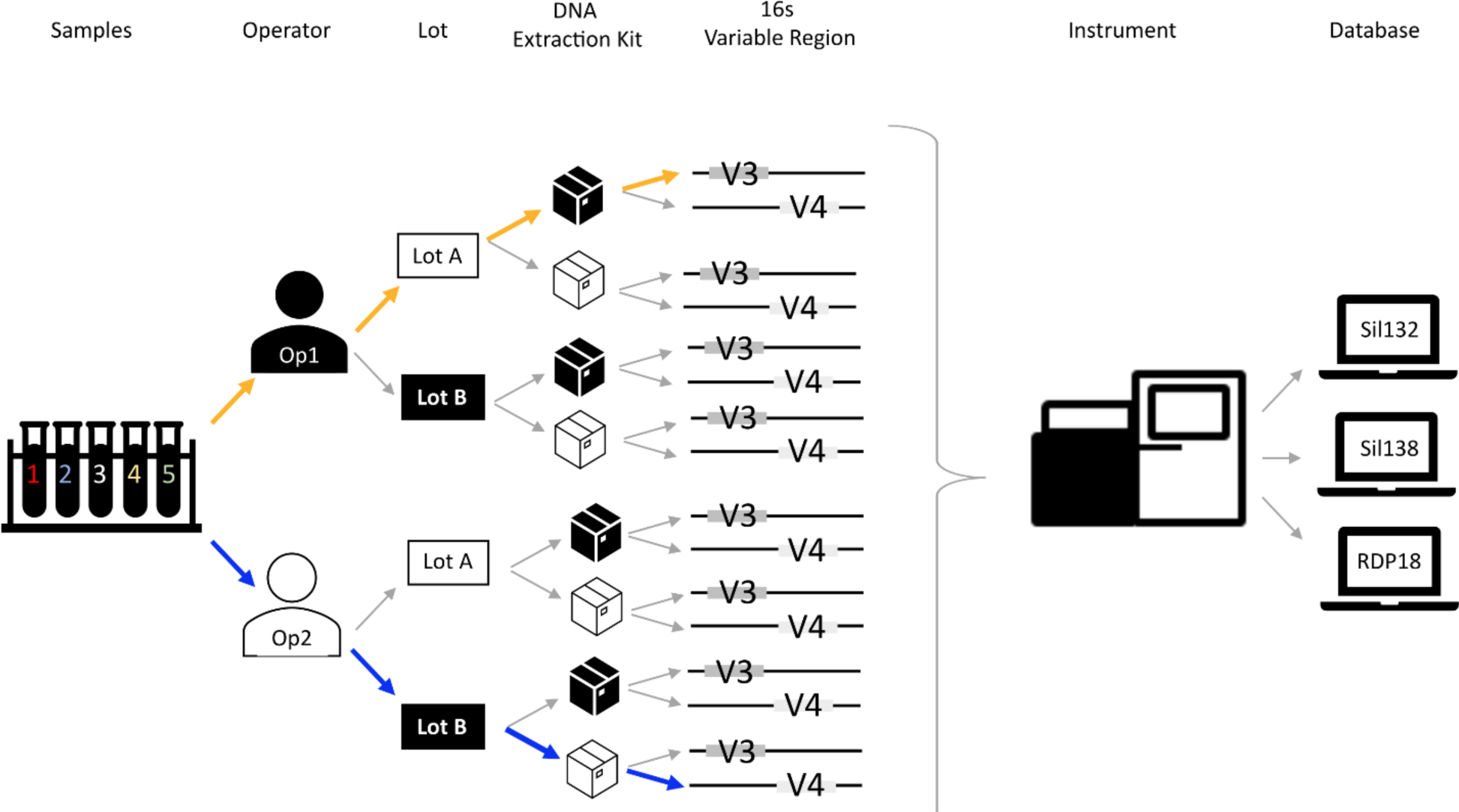
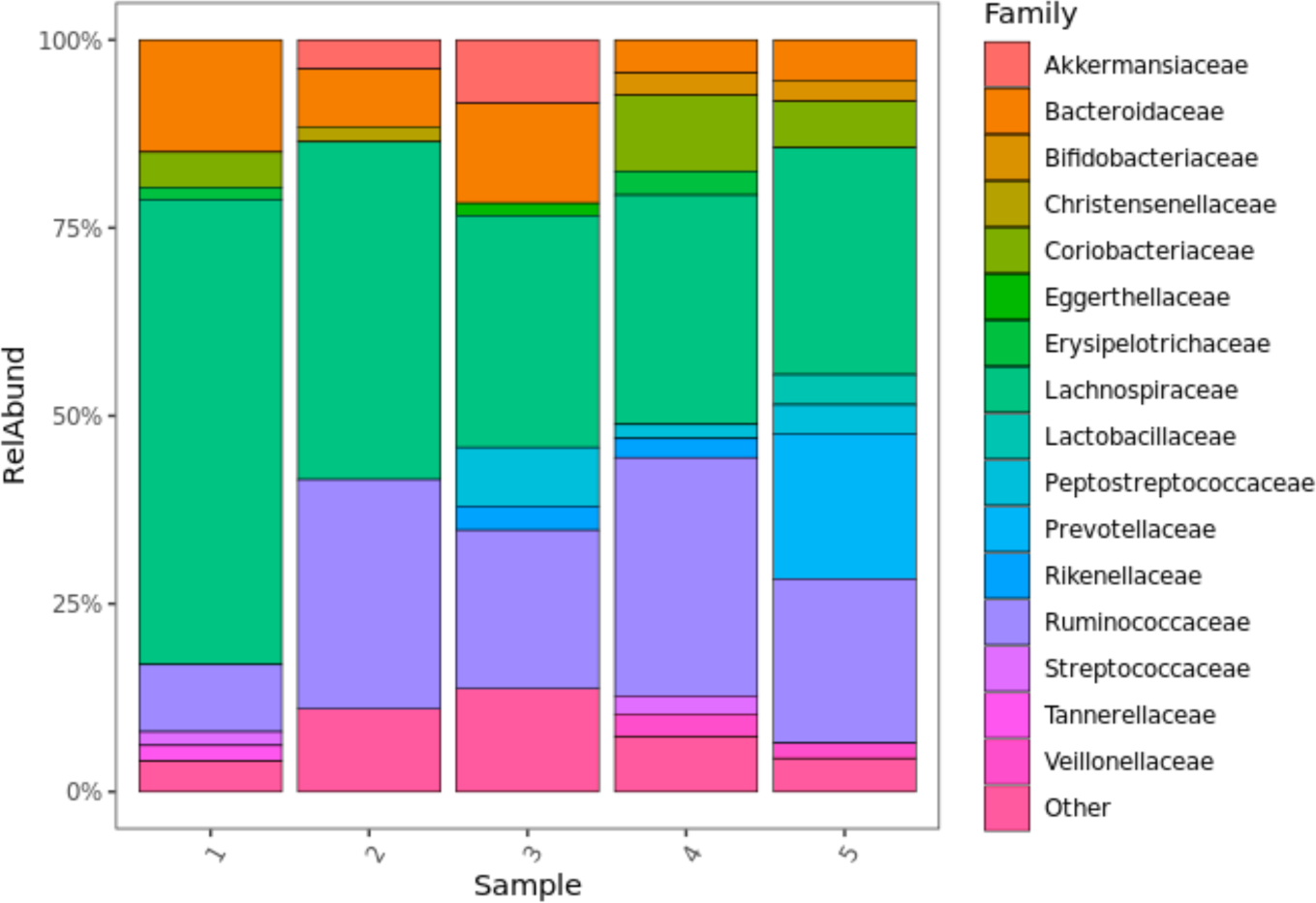
Systematic experimental design enables quantitative bias assessment. The graphic above illustrates the levels compared for each step of the metagenomic workflow. In total, 5 different human stool samples were handled by 2 operators, preparing samples from 2 lots, for MGS analysis using 2 DNA extraction kits and 2 variable regions, with taxonomic assignment coming from 3 databases. The top (orange) and bottom (blue) arms of the experimental design represent completely orthogonal protocol choices through all wet-bench operations, and these two protocols were replicated (2x) for each sample and database to explore reproducibility. This resulted in a total of 240 MGS data files (270 with replicates) that were analyzed.

This work represents a proof-of-concept and was intended to demonstrate a full factorial analysis of a metagenomic sequencing workflow. Ideally, the implementation of a full-factorial experimental design would specify ‘high’ and ‘low’ values for each selected parameter to bound the range of normal experimental conditions. In the case of the metagenomics workflows evaluated here, the selected experimental parameters of interest exhibited discrete, non-numerical, non-continuous levels (e.g., Operator, Lot, Extraction Kit), and the selected options (levels) may not bound the normal range of experimental conditions. Herein, a rigorous statistical approach was demonstrated for evaluating bias in metagenomics workflows that others could utilize in their own laboratories with their particular protocols.

### MGS: Compositional Analysis

A typical metagenomic analysis may include bar charts to show relative abundance and community members. As such, the composition of each sample for a single workflow was visualized in a relative abundance bar chart at the Genus taxonomic level (Figure 3A). The differences between stool samples were evident, and good reproducibility was observed between replicate analyses. For example, while *Lachnospiraceae* and *Ruminococcaceae* make up the plurality of each sample, their relative proportions vary, and some taxa (e.g., *Rikenellaceae, Prevotellaceae*, *Akkermansiaceae*) appeared variably present, occurring above the 1.5 % cutoff in some samples but not others. A comparison of MGS bar charts from alternate workflows showed that differences in sample composition could be correlated with methodological choices (Figure SI-1); however, using bar charts to assess methodological parameters can be difficult to implement systematically. While differences between samples can be readily observed, it is difficult to determine the magnitude of these differences or to correlate them with particular methodological decisions. Also, comparing the compositional changes between many individual bar charts proves to be cumbersome and challenging to automate.

**Figure 3:**
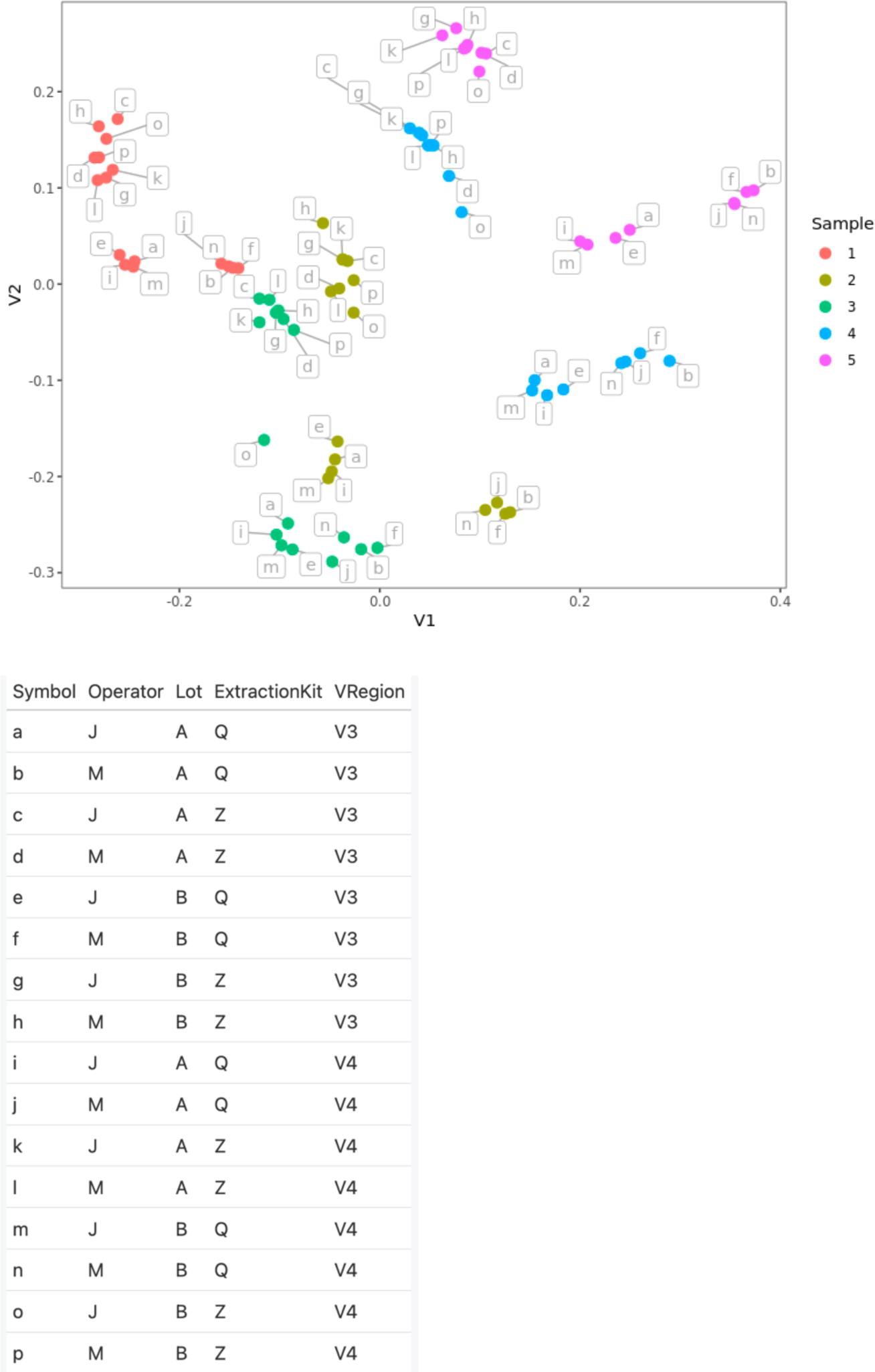
(a) Bar charts and (b) PcoA plots from 5 human stool samples. (3A) Family level relative abundance bar charts show the compositional diversity among 5 human stool samples for a single workflow. For this plot, Families present at <1.5% relative abundance are grouped as “Other”. Correlation between the observed compositions and specified protocol choices (see Figure SI-1) can help identify particularly sensitive steps in an MGS workflow. (3B) Bray-Curtis dissimilarities can also be used to identify sensitive MGS protocol choices. Points are colored here to indicate fecal sample and letter flags show additional protocol choices for each dataset, as denoted in the table. Further clustering (see Figure SI-2) can be used to rank protocol choices with respect to their impact on the observed composition. Though these two plots are conventionally used, they are limited in their utility to identify and address biases.

Another common approach is to evaluate shifts in composition by calculating similarities (or dissimilarities such as Bray-Curtis) between MGS datasets and then generating a principal coordinate analysis (PCoA) plot from the resulting distance metrics (Figure 3B). In this approach, a single point was generated for each unique combination of sample and analysis workflow, and clusters of points were then correlated with methodological parameters. For example, with the data plotted in Figure 3B, data from a common sample (same colors) generally clustered, with some overlap and sub-clustering associated with methodological parameters (denoted with letter flags). To compare between specific methodological choices, additional faceting generally allowed for their relative impact to be assessed (Figure SI-2a-e). This pattern recognition approach facilitated the evaluation of multiple datasets simultaneously; however, the results remained mostly qualitative in nature, as the apparent significance of each evaluated parameter depended on the ordination conditions of that dataset.

### MGS: Ratiometric Analysis

An additional challenge when comparing MGS results between samples is the compositional nature of the datasets. That is, the measured relative abundance value for any taxa depended on both its actual abundance in the sample and on the sum of the observed abundances for all other taxa present in that sample. An alternative approach for comparing MGS results has focused on ratiometric comparisons, and these ratios of taxa have been shown to be *independent* of sample composition. (5, 8, 16-18) Whereas the initial relative abundance values could be misleading, ratios of relative abundance between two taxa within a sample have been shown to be more reliable, varying with just the actual taxa abundances and measurement biases.(8)

This concept has been implemented previously with native taxa, such as calculating the *Firmicutes:Bacteroidetes* (F:B) ratios (among others), with varying levels of success (10, 19-21). To demonstrate ratiometric analysis, we examined the F:B ratio within our experimental design (Figure 4). The F:B ratio provided a single calculated value for each unique combination of stool sample and experimental workflows (black points), with the samples analyzed under identical workflows connected by light grey lines. For reference, the two completely orthogonal workflows are colored orange and blue (and match the top and bottom arms of the experimental design in Figure 2).

**Figure 4:**
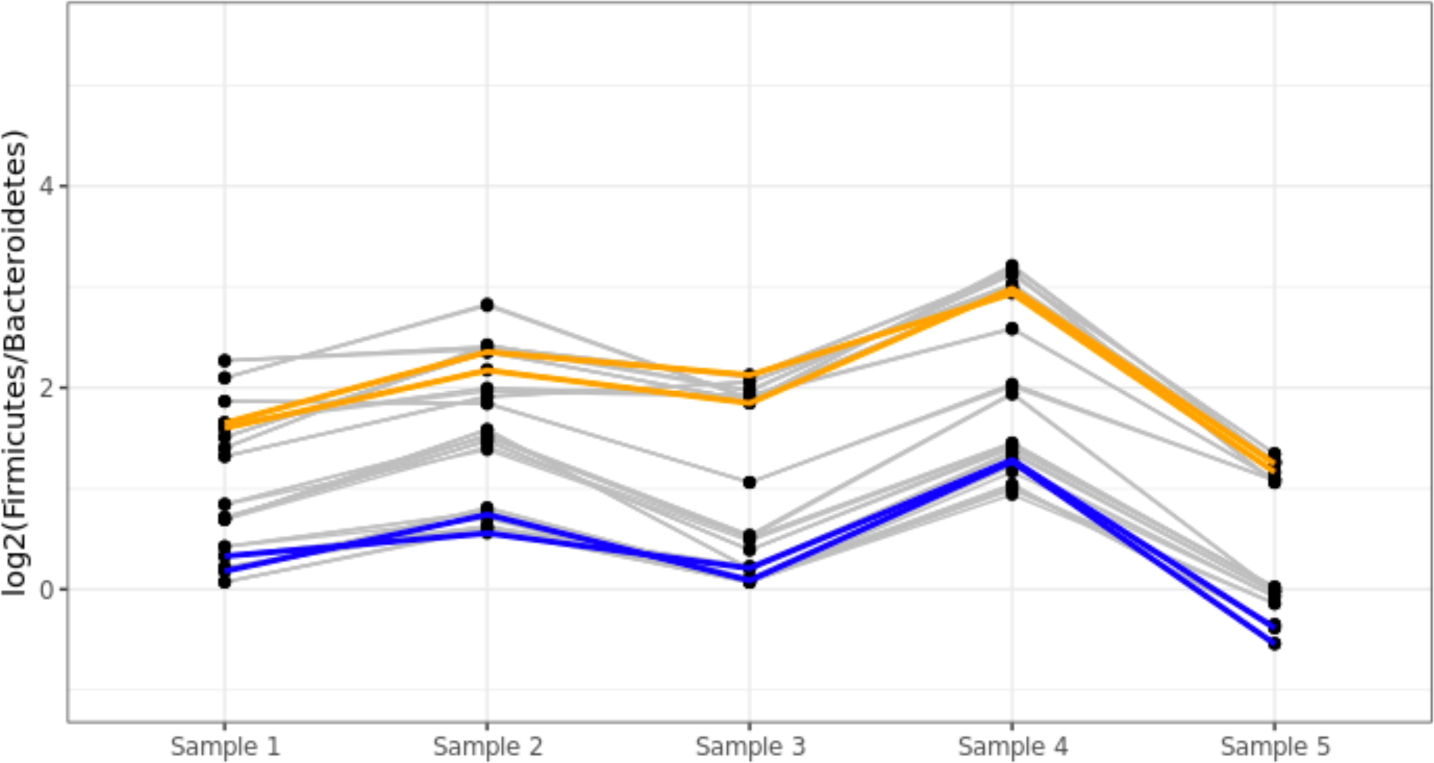
Impact of methodological parameters on the Firmicutes:Bacteroidetes ratio. The Firmicutes:Bacteroidetes ratio was determined for each sample (black points) under every combination of the methodological parameters investigated in this study (48 combinations for each of the 5 samples). A grey line connects the dots between fecal samples processed using the same protocol; dots and lines overlapped for protocols where the varied parameters didn’t change the observed Firmicutes:Bacteroidetes ratio. The blue and orange lines denote the results from the top and bottom legs of our experimental design flowchart (shown in Figure 2), respectively, and represent orthogonal choices for every methodological parameter (operator, lot, DNA extraction kit, variable region, database). There are two blue and two orange lines, as each of these workflows were run in duplicate to demonstrate reproducibility when all parameters are held consistent.

As expected, due to real variation in the actual F:B ratio between biologically distinct samples, similar trends (‘M’ shape) were observed across samples for each of the different protocols evaluated. A Kruskal-Wallis Test confirmed that there was a statistically significant difference in F:B ratio between the stool samples (χ^2^(4) = 77.8, p = 1e-15). However, within each sample, a greater than 4-fold difference in F:B ratio was observed between different workflows, reflecting the impact of methodological choices on the observed F:B ratio. Indeed, this workflow-dependent effect (bias) was at least as significant as the biological differences observed between distinct stool samples. In a rapidly evolving field like MGS, this kind of workflow-dependent measurement bias significantly complicates comparisons between studies.

### MGS: Internal Standards

Given that native taxa are expected to vary naturally between the stool samples, as observed here with Firmicutes and Bacteroidetes, an alternative approach for ratiometric analysis is to add exogenous strains (spike-ins) to serve as internal controls. In this study, two strains were uniformly added during the batch production of these stool samples. *Leifsonia xyli* (Gram positive) and *Aliivibrio fischeri* (formerly known as *Vibrio fischeri*, Gram negative), a plant pathogen and marine organism, respectively, were selected as they are unlikely to be found in human stool samples and could therefore be uniquely identified taxonomically. Unlike the native taxa, these spike-in organisms were added at consistent abundances across all stool samples.

Similar to the F:B ratio plotted in Figure 4, the *Leifsonia*:*Aliivibrio* (L:A) ratio was calculated for each unique combination of stool sample and experimental protocol (Figure 5). As before, a high degree of bias was observed between individual workflows (vertical distribution of points for each sample). However, unlike the previous analysis, the L:A ratio did not exhibit trends across samples (‘M’ shape from Figure 4). Supporting this lack of biological variability, a Kruskal-Wallis Test failed to find a statistically significant difference in L:A ratio between the stool samples (χ^2^(4) = 4.8, p = 0.31). Instead, observed variations in the L:A ratio between samples analyzed with a common workflow (i.e., following a single line trace) were attributed to measurement variability. Considering the differences in the relative abundances of *Firmicutes* and *Bacteroidetes* (≈88 %) versus *Leifsonia* and *Aliivibrio* (≈0.6 %), the increased measurement variability observed in Figure 4 was unsurprising. Since the spike-in organism abundances were detected consistently between samples, we concluded that either of these spike in organisms were suitable for use as internal standards for ratiometric analyses. For the purposes of demonstrating ratiometric analysis we chose to use *Leifsonia*. As such, comparing the *Bacteroidetes:Leifsoni*a ratio between different samples handled under a constant workflow allowed actual changes in Bacteroidetes to be quantified independent of any other native taxa.

**Figure 5:**
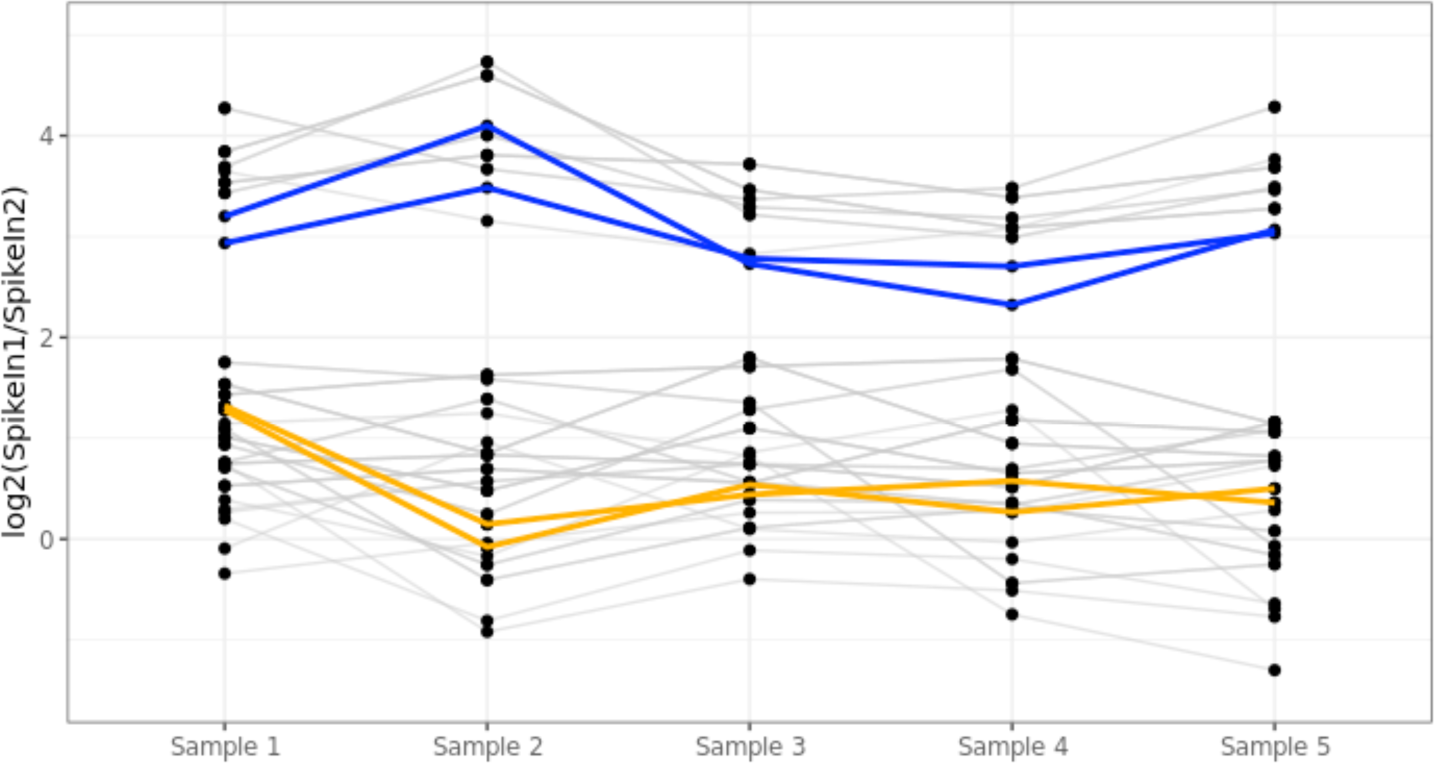
Analysis of internal standards added uniformly to all stool samples. Two exogenous organisms, Aliivibrio and Leifsonia, were added as internal standards to the stool samples. The ratio of Aliivibrio:Leifsonia was calculated for each sample (black points) under every combination of the methodological parameters investigated in this study (48 combinations), as described in Figure 4. A grey line connects the dots between fecal samples processed using the same protocol; dots and lines overlapped for protocols where the varied parameters didn’t change the observed Aliivibrio:Leifsonia ratio. The blue and orange lines denote the results from the top and bottom legs of our experimental design flowchart (shown in Figure 2), respectively, and represent orthogonal choices for every methodological parameter (operator, lot, DNA extraction kit, variable region, database). There are two blue and two orange lines, as each of these workflows were run in duplicate to demonstrate reproducibility when all parameters are held consistent.

### Quantifying Parameter Effects

In Figure 5, the systematically varied workflows resulted in significant variation in the ratiometric MGS results within each sample. The full-factorial experimental design (Figure 2) was structured to balance each decision point in the workflow and facilitate independent characterization of each methodological choice. A Parameter Effect was calculated for each evaluated step in the workflow (i.e., parameter) and the method selected (i.e., level) by calculating the fold change of ratiometric results from workflows using the specified method versus the average results across all workflows (Equation 1)

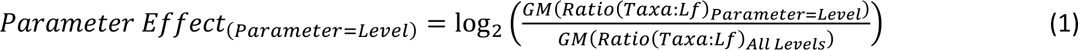

where *GM* denotes the geometric mean. This calculated parameter effect allowed individual workflow steps and specific methodological choices to be directly compared.(11, 12)

As expected, when the Parameter Effects for the ratio of the two internal standard taxa were calculated, (Figure SI-3) no significant effect was measured between samples due to the even amount of the internal standards added to each sample; however, several of the methodological parameters did exhibit statistically significant Parameter Effects for the ratio of the two internal standards. It was important to note that the Parameter Effect calculation (Equation 1) was based on ratios with the internal standard and could not attribute observed effects to either of the ratioed taxa individually. When we subsequently considered taxa native to the stool, a large Parameter Effect denoted a large bias differential between the taxa-of-interest and the internal standard, while a small or negligible Parameter Effect indicated a similar response between the taxa-of-interest and the internal standard.

The Parameter Effect calculations for the phyla Bacteroidetes revealed significant effects for several parameters (Figure 6), additionally the differences between biological samples (e.g. aliquots) were also generally significant for Bacteroidetes. One benefit of the Parameter Effect calculation in the context of a full-factorial design was its ability to quantify the effects of methodological parameters even in the context of biological variations. This was confirmed by calculating the Parameter Effects for *Bacteroidetes* within each stool sample individually (Figure SI-4). The within-sample Parameter Effects matched the aggregate Parameter Effects, albeit with a larger (less precise) confidence interval due to the smaller number of samples. In the aggregate Parameter Effects calculation, the extraction kit and operator parameters exhibited statistically significant effects, while the parameters of lot, 16S variable region, and database did not exhibit significant effects.

**Figure 6:**
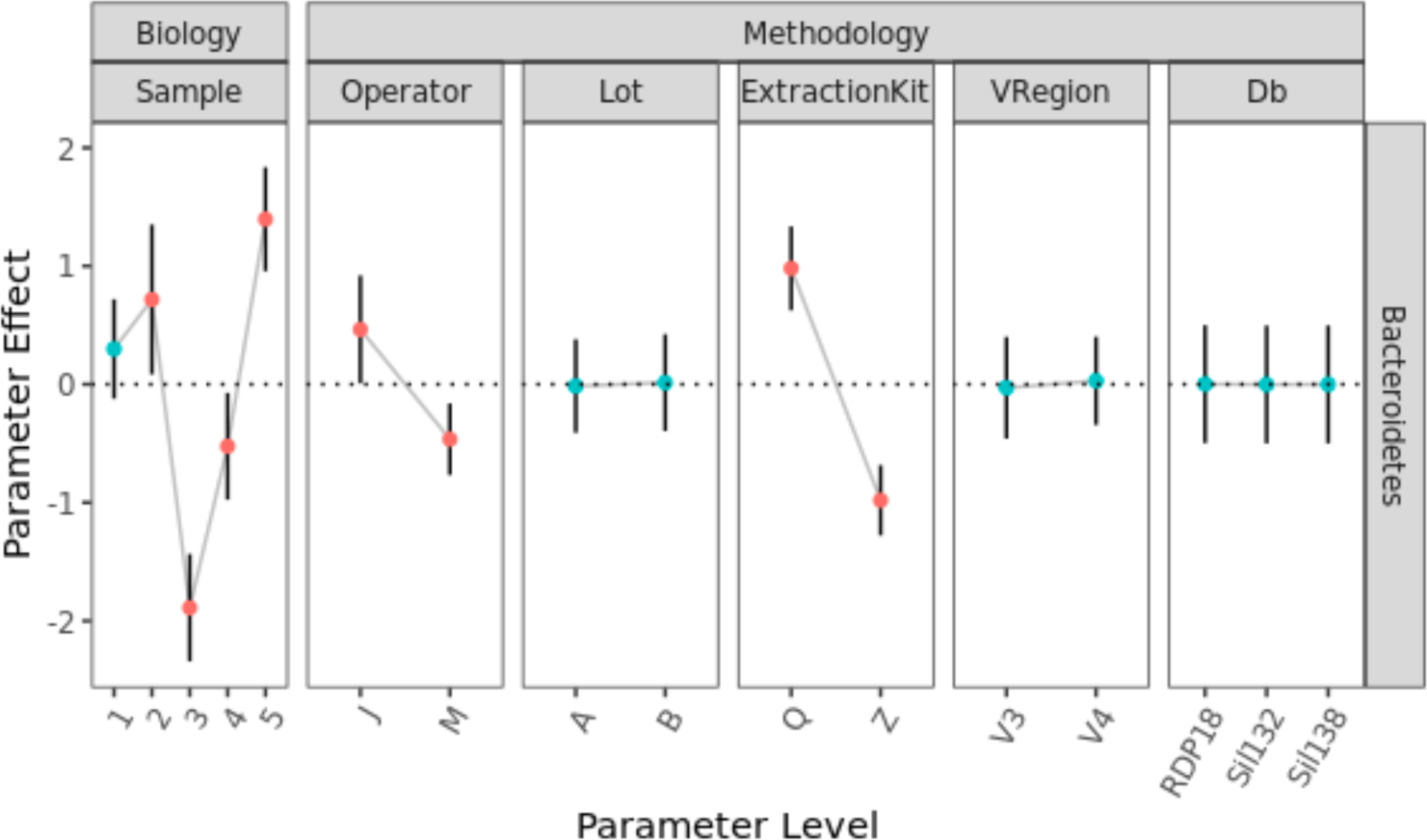
Quantitative comparison between methodological parameters. The Bacteroidetes:Leifsonia ratio was calculated for each dataset to enable comparison between protocols. The effect of each parameter was calculated by dividing the average ratio for all datasets at the denoted parameter level by the average ratio across all parameter levels, as shown in Equation 1. This parameter effect was plotted as a fold change on a log2 scale, such that the horizontal line at 0 denotes the null hypothesis of no effect. The magnitude of the effect of protocol choices (e.g., Extraction Kit) can be directly compared between parameters and parameter levels. Data error bars showed 99 % confidence intervals, and the points that are statistically significantly different from the mean (p<0.01) were colored red.

The Parameter Effects for Extraction Kit were similar in magnitude to the real biological differences between samples, potentially complicating comparisons between samples that were analyzed using different methods. For example, comparing stool samples 4 and 5 using Extraction Kit Z, while holding all other methods constant, revealed an actual difference in *Bacteroidetes:Leifsonia* ratio (B:L) of 4.1-fold. However, evaluating only sample 4 using Extraction Kits Q and Z revealed an apparent difference in B:L of 4.4-fold, just due to the methodological difference. When the measured B:L was compared between an analysis of stool sample 4 using extraction kit Q and an analysis of stool sample 5 using extraction kit Z, the results were essentially identical (1.05-fold difference) because the effects of the biological and methodological differences were of similar magnitude and in opposite directions. Alternately, analyzing sample 4 with extraction kit Z and sample 5 with extraction kit Q yielded an apparent change in B:L of 10.3-fold because the biological and methodological differences compounded.

In addition to Bacteroidetes, Parameter Effects were calculated for multiple independent phyla across the 5 stool samples (Figure SI-5). As expected for diverse stool samples, the native phyla varied in different ways between the five fecal samples, reflecting their disparate biological compositions. However, the calculated Parameter Effects for methodological parameters also varied significantly between phyla. For instance, Proteobacteria and Bacteroidetes exhibited significant Parameter Effects for Extraction Kit, while Actinobacteria and Firmicutes did not exhibit significant Parameter Effects for Extraction Kit (Figure SI-5). It was notable (while not unexpected) that these calculated Parameter Effects were taxa specific.

### Quantifying Parameter Effects: Selected genera

While the analyses of native microbes presented thus far have focused on the highest taxonomic classification (i.e., phyla), much published research has linked beneficial or deleterious biological outcomes with more specific taxonomic clades.(22) Thus, we calculated Parameter Effects for a panel of genera previously posited as having particular interest for gut health and representing multiple genera within specified phyla (Figure 7). Overall, the methodological factors of lot and variable region did not exhibit significant Parameter Effects for any of the genera evaluated here (and have been omitted from the plot), while the factors: Operator, Extraction Kit and Database were sometimes significant.

**Figure 7:**
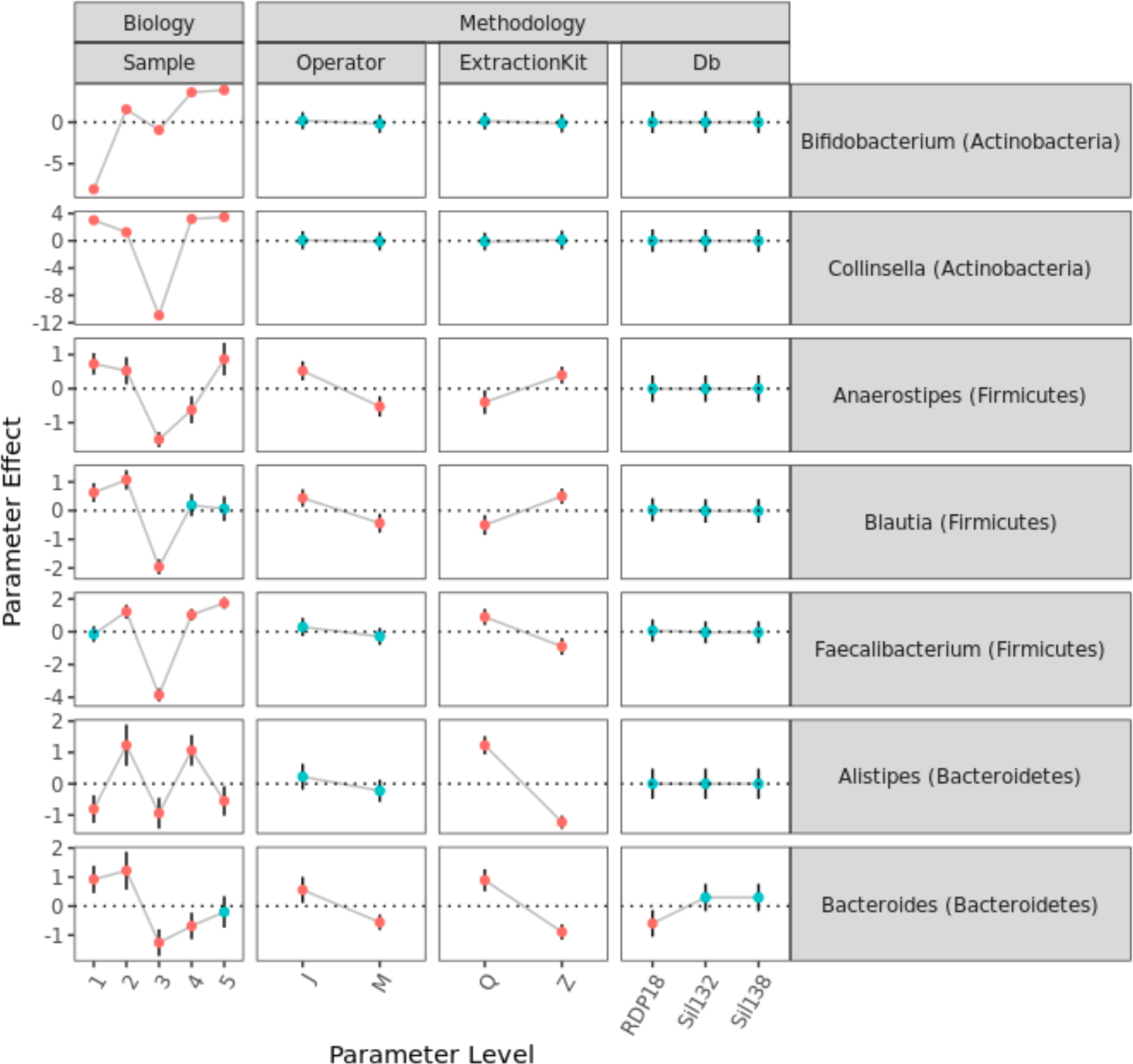
The effect of methodological parameter choices impacts the quantitation of some Genera associated in the literature with the health of the gut microbiome. Like Figure 6, parameter effect was calculated for the ratio of various taxa of interest to an internal control (Leifsonia). The parameter effect was calculated by dividing the average results for the specified parameter level for the specified ratio by the average results across all factor levels for the specified ratio. Data error bars show a 99 % confidence interval, and statistically significant points (p < 0.01) are colored red. The effects broken out by taxonomic levels are plotted in SI Figures 6a-6g.

Interestingly, while the two genera within the *Actinobacteria* phyla exhibited similar responses to the methodologies evaluated in this study, this was not the case within the *Firmicutes* or *Bacteroidetes* phyla. Within the *Firmicutes*, the *Faecalibacterium* genus was preferentially enriched by one extraction kit and exhibited no Parameter Effect for operator, while the *Blautia* and *Anaerostipes* genera were preferentially enriched by the other extraction kit and exhibited significant Parameter Effects for operator. Within *Bacteroidetes*, the *Alistipes* and *Bacteroides* genera exhibited similar Parameter Effects for extraction kit, but differed in the Parameter Effects for operator and database. These variations in the Parameter Effects within Phyla demonstrated significant phenotypic heterogeneity at the genus level and underscored the difficulty in identifying reference organisms as potential surrogates for other members within a taxonomic grouping. To further demonstrate the variation in Parameter Effect an expanded view of each taxonomic level (Phylum, Class, Order, Family) for the genera in Figure 7 is present in Figure S1-6. In some cases, trends stayed the same as you move down the taxonomic hierarchy (Figure SI-6e, operator) while in others significant differences were only observed at high taxonomic levels (Figure SI-6f, operator) or low taxonomic levels (Figure SI-6f, Extraction Kit). The scope of the current effort was limited to the specific parameters varied and the parameter levels employed; however, similar conclusions (i.e., taxa-specific Parameter Effects that can vary substantially within taxonomic groupings) should be expected for other MGS protocols.

### Quantifying Parameter Effects: All Taxa

Using ratios with the internal standard, we calculated Parameter Effects for each individual taxa native to the fecal samples. The methodological parameter which generally exhibited the most significant effect was Extraction Kit, and we focused on that Parameter Effect to assess the degree of variation between taxa (Figure 8). Additionally, discrepancies in taxa names between databases or even database versions made systematic comparisons of individual taxa challenging, so this quantification of the Parameter Effect for extraction kit was performed within the Silva 138 database.

**Figure 8:**
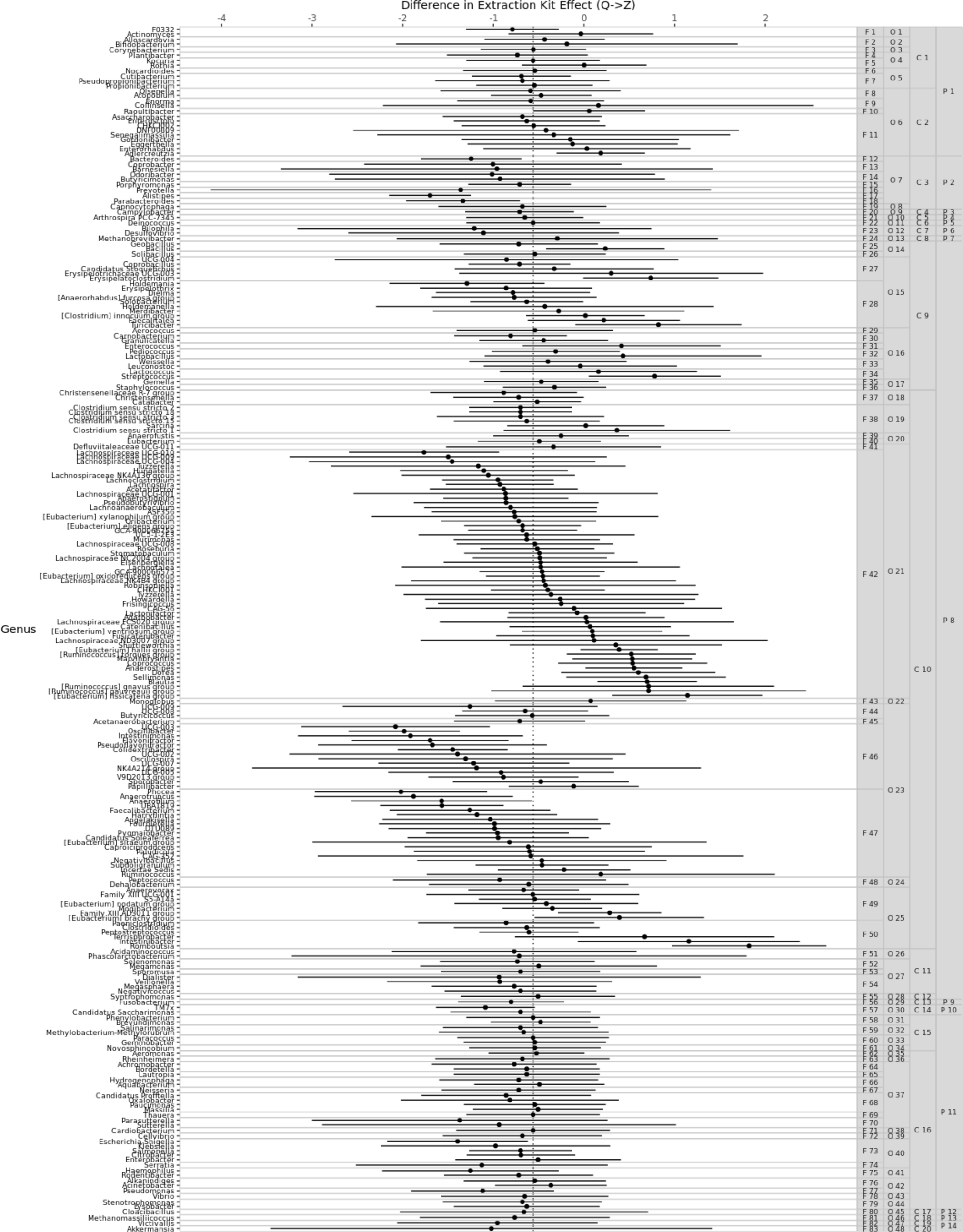
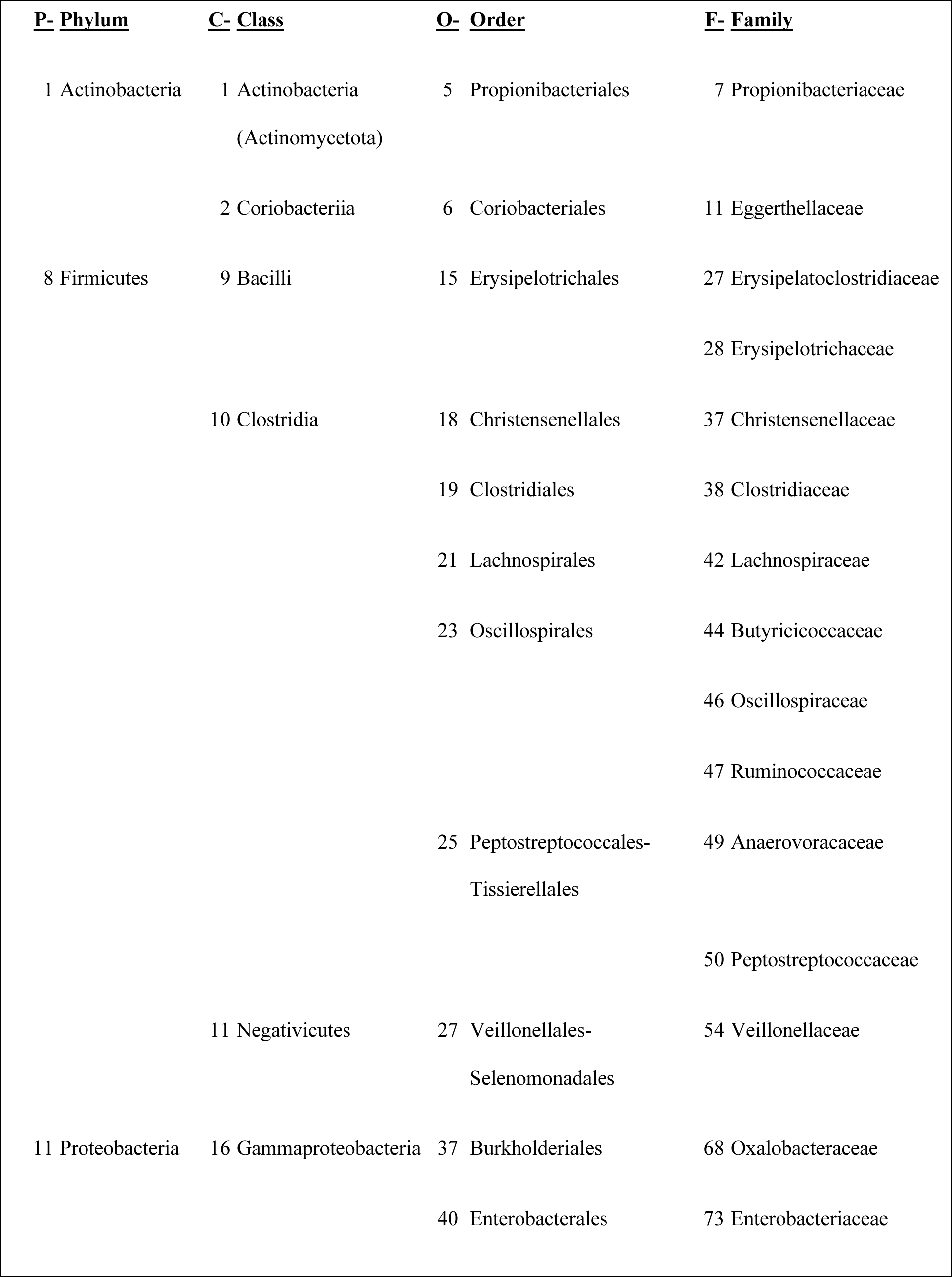
The Effect of Extraction Kit on Different Genera. For each genera in the fecal samples, the difference in the Parameter Effects of the two Extraction Kits (*Effect_Kit=Z_* – *Effect_Kit=Q_*) was calculated. (Leifsonia was used as the internal standard.) Error bars show a 99 % confidence interval. Genus name is given on the left hand of the figure, with corresponding Phylum, class, order, and family groupings on the right. The table below the figure provides the taxonomic key for all Families with ≥3 Genera, and the full taxonomic key (249 genera) is included in the Supporting Information (SI Table 1).

For some taxa, such as those in the family Clostridiaceae (Figure 8, F38), there appeared to be little or no effect of extraction kit, with all 99% confidence intervals overlapping with the mean and no statistical difference between the 6 genera. In comparison, the Peptostreptococcaceae family (Figure 8, F50) exhibits a wide range of extraction kit effects (and statistically significant differences) between the 6 genera. In general, families with a greater number of genera present in the sample showed a greater range of extraction kit effects. While this observation was not surprising, it was notable that even closely related taxa could be impacted differently by methodological choices such as extraction kit. Figure 8 would certainly look different in the details for other methodological choices or MGS workflows, but this demonstrated that in general the effects of methodological parameters (i.e, method bias) could vary substantially between taxa, even closely-related taxa, and cast doubt on the ability of well-characterized ‘reference species’ to serve as adequate surrogates.(23) This observed lack of consistency at the genera level has important implications for developing mock communities that will adequately stress a workflow (24). Instead, these data suggested that method bias for particular taxa-of-interest should be measured through experimental characterization of the protocols being considered.

### Correcting for Methodological Bias

Mathematically, quantification of Parameter Effect provided a mechanism for achieving comparability between differing protocols.(8) The observed differences between protocols in the observed relative abundances of individual taxa (i.e., method bias) could be computationally brought into agreement by correcting for the taxa-specific Parameter Effect. To validate this experimentally, Genus-level Parameter Effects were calculated for the methodological parameters of extraction kit and operator. These two parameters were selected because they had previously exhibited significant Parameter Effects for multiple tested Genera. Additionally, the Parameter Effects were only calculated using data from a single stool sample to simulate the ability of this approach to improve MGS characterization of previously untested samples. Stool sample 3 was used to calculate the Parameter Effects because it exhibited the most unique Genera. Using the calculated Parameter Effects, the observed abundances of Genera in all five samples were then computationally corrected to account for the protocol employed (i.e., which extraction kit and operator were specified). The initial- and corrected-relative abundances were used to calculated Bray-Curtis Dissimilarities between samples across all protocols as seen in the principal coordinate analysis plot colored by Sample, with the variability between protocols approximated with a 95 % data ellipse (Figure 9). The effect of the computational correction was evident in the reduced size of the data ellipses for each sample, demonstrating a significant decrease in the observed dissimilarities between data collected under differing protocols.

**Figure 9:**
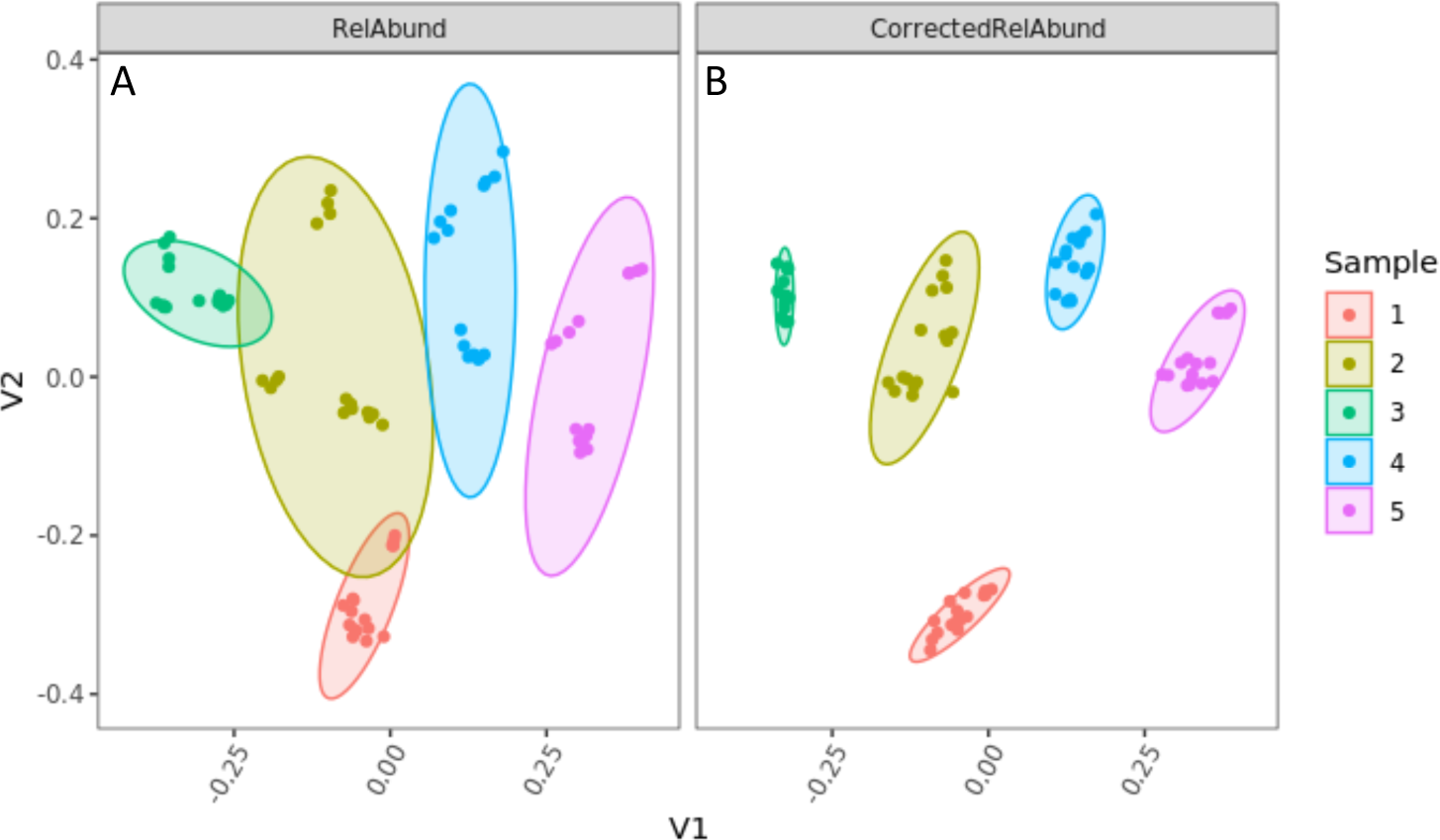
Correcting for Extraction Kit and Operator Biases. Genus level Parameter Effects were calculated for Extraction Kit and Operator using data stool sample 3, and the Relative Abundances of individual Genera in all five samples were computationally corrected to account for the protocol used (i.e., which extraction kit and operator were specified). Bray-Curtis dissimilarities were calculated between all datasets and displayed on principal coordinate plots of (A) Relative Abundance or (B) corrected Relative Abundance, colored by fecal sample. 95 % data ellipses are shown to highlight the amount of dispersion observed between differing protocols for both conditions.

It was notable that the improved comparability (reduced method bias) between protocols was observed for all five stool samples, even samples 1, 2, 4, and 5 that were not included in the Parameter Effect calculations. The ellipse centroids did not shift with computational correction because the Parameter Effect calculation was centered around a fold-change of 0 (no effect), and the arms of the experimental design were evenly balanced. It should also be noted that this computational correction did not necessarily improve the resulting data with respect to the actual biological composition of the samples. Rather, the data have been made more comparable between protocols by accounting for their taxa-specific biases. In this way, the dissimilarities within the datasets for each sample were reduced. This result strongly supported the idea that MGS data collected under differing protocols could be harmonized through the systematic analysis of common samples (e.g., reference materials).

## Conclusions

The impact of methodological variables on MGS results is generally well recognized. In the current manuscript we combined a full factorial experimental design strategy with ratiometric analyses to quantify the impact of specific methodological choices. When the F:B ratio was calculated for individual stool samples, a >4-fold divergence in results was observed between different protocols (Figure 4). Replicate analyses confirmed that MGS results were reproducible run-to-run, so the observed variations were attributed to bias associated with specific methodological differences in the analysis protocols. Furthermore, this methodological bias was similar in magnitude to the observed differences between the biologically distinct samples.

Using the full factorial design, the Parameter Effect could be calculated to directly compare between specific biological and methodological parameters (Figure 6). Among the 5 methodological parameters evaluated in the current investigation, extraction kit and operator were statistically significant, while stool lot, 16S variable region, and database were not statistically significant. The observation that extraction kit introduced significant measurement bias was not novel; however, the benefit of the full-factorial experimental design was that the Parameter Effect for each variable could be quantified and directly compared to the Parameter Effect associated with biological variations. For example, for extraction kits specifically, the observed Bacteroidetes measurement bias was sufficiently large to mask real differences between distinct biological samples. This observation raises questions about attempts to compare MGS data collected under different protocols. Somewhat surprisingly, operator also yielded a Parameter Effect that was weak but statistically significant and showed that human variability can still impact results even with locked-down protocols and identical lab equipment.

By combining the full factorial design with ratiometric analysis against internal control organisms, Parameter Effects could be calculated for individual taxa-of-interest native to the stool samples. For a panel of 7 genera posited in the literature to be associated with specific gut-health outcomes, we observed a large diversity of Parameter Effects (Figure 7), with some taxa exhibiting minimal measurement bias for the methods evaluated here, while several others exhibiting significant bias across multiple methodological decisions (Figure 7). Notably, Parameter Effects did not appear to group by taxonomic clade (e.g., phylum) and this somewhat unexpected finding warrants further investigation. Focusing specifically on the extraction kit, some related taxa exhibited similar Parameter Effects while other closely related taxa (e.g., genera within a family) exhibited quite divergent Parameter Effects. This diversity of methodological bias could have important implication for reference material development (i.e., mock communities) where microbial mixtures often contain only 10s of strains. These findings suggest that the generalizability of method performance based on a small subset of strains could be very limited and justifies efforts to develop more complex mock communities and natural surrogate reference materials that contain many more taxa and are compositionally similar to samples of interest. Indeed, using genera-specific Parameter Effects calculated for one of the samples investigated here, method bias between divergent protocols was significantly reduced for all samples.

The current report details some of the measurement challenges within the metagenomic space and presents a method to systematically assess sources of bias. However, there are a few limitations worth noting. As designed, this study only examined a small subset of methodological variables, while holding all others constant. This enabled the full-factorial experimental design and allowed Parameter Effects to be quantified for those variables. This study was not meant to serve as a comprehensive review of available kits, library preparations, NGS technology, databases, etc. largely due to a second limitation of the full factorial design: it is resource intensive and expensive. A significant amount of starting material was needed to ensure a homogenous sample supply, a notable challenge for clinical and archival samples that are routinely limited. As reagent costs and personnel time requirements quickly scale in increasing numbers of methodological variables, a partial-factorial experimental design may be sufficient in many cases. Nevertheless, as technology advances, the quantitative determination of the bias associated with specific changes to the workflow can make future results backward compatible.

In summary, we presented an experimental design and analysis strategy that enables direct quantitative comparison of different biological and methodological parameters. We encourage MGS research groups to consider characterizing the impact of their in-house protocol choices on their MGS results. As more labs seek to improve the comparability of their MGS results between divergent protocols, this approach provides a quantitative strategy for assessing methodological bias and shows how data collected under differing protocols can be harmonized by widespread analysis of a common sample (e.g., reference material).

## Methods (Mosaic_Materials-and-methods.docx)

### Samples and Sample Handling

Samples used in this study were previously described.(15) Briefly, five samples were generated from five different donors. Each sample was prepared by pooling and homogenizing 4 consecutive bowel movements from the donor, combining homogenized material with Omnigene Gut Solution, spiking with 10^8^ CFU/mL of *Aliivibrio-fischeri* (Gram negative, formerly known as *Vibrio fischeri*) and *Leifsonia xyli* (Gram positive), and then aliquoting 1 mL aliquots. The final concentration of the material was 100mg/mL. Samples were received as a set of five, one sample from each donor (labeled 1-5). For this study we ordered two sets (designated Lot A and Lot B) of the samples. Samples were stored at −80 °C until the experiments were conducted. On the day of the experiment, samples were vortexed vigorously and then 100 µL from each sample (1–5) was aliquoted into five 1.5-mL microcentrifuge tubes. This process was repeated for both lots. Operator 1 used three aliquots of each sample (1–5) from Lot A, and two aliquots from each sample (1–5) from Lot B. Operator 2 used two aliquots of each sample (1–5) from Lot A, and three aliquots from each sample (1–5) from Lot B.

### DNA Extraction

#### Genomic DNA Extraction Kit Q

Samples were extracted using two different methods. Method 1 was done with the QIAamp Fast DNA Stool Mini DNA prep protocol (Qiagen, Cat# 51504). Briefly, 1 mL of the Inhibit EX buffer was added to each of the samples and vortexed (Mo Bio Vortex – Genie 2 at max speed) for 1 minute. The samples were then centrifuged (Eppendorf Centrifuge 5417R) at 16,000×*g* for 1 minute to pellet the samples. A new 2 mL centrifuge tube was prepared containing 25 μL of proteinase K and 600 μL of the supernatant from the original tube was added to the tube containing proteinase K (supplied with kit). This was followed by the addition of 600 μL of Buffer AL; samples were vortexed for 15 seconds to mix. The samples were incubated for 10 minutes and 70 °C. After incubation, 600 μL of molecular biology grade ethanol (100 %, Sigma-Aldrich, E7023-1L) was added and vortexed. In 600 μL increments, the lysate was loaded onto the QIAamp spin column, and samples were then centrifuged 16,000×*g* for 1 minute. After centrifugation, columns were placed in new 2 mL collection tube and the flow-through was discarded. This process was repeated until all the lysate had been loaded onto the column. Next, the columns were washed first with 500 μL of Buffer AW1 (centrifuged 16,000×*g* for 1 minute), then 500 μL of Buffer AW2 (centrifuged at 16,000×*g* for 3 minutes). Flow-through was discarded and tubes replaced between each wash. To remove any remaining wash solution, columns were placed in a new collection tube and centrifuged at 16,000×*g* for 3 minutes. Columns were then placed in 1.5 mL labeled microcentrifuge tubes and 100 μL of Buffer ATE was added to the center of the membrane. The samples incubated for 5 minutes and then centrifuged at 16,000×*g* for 1 minute to elute the DNA.

#### Genomic DNA Extraction Kit Z

Method 2 was done with the ZR Fecal DNA miniprep (cat#D6010) from Zymo Research following the manufacturer protocol with minor modifications. Briefly, each sample was combined with lysis buffer and loaded onto the MoBio vortex Genie 2 with 2 mL tube adapter for 20 minutes at full speed. The lysis tube was centrifuged at 10,000×*g* for 1 minute. Then 400 μL of the lysate was added to the IV spin filter and centrifuged at 7,000×*g* for 1 minute. The process was repeated until all lysate had been passed through the spin filter. Filtrate was combined with 1,200 μL of the fecal DNA binding buffer and mixed well by pipetting. The mixture was transferred to the IIC column in 800 μL increments and centrifuged at 10,000×*g* for 1 minute. The supernatant was discarded. The bound DNA was washed and eluted in 150 µL of the elution buffer after incubation at room temperature for five minutes. The DNA was then transferred to the prepared IV-HRC spin filter and centrifuged at 8,000×*g* for three minutes.

Extracted DNA from both methods was quantified using DeNovix dsDNA High Sensitivity (Catalog number: KIT-DSDNA-HIGH-2) and analyzed on the DeNovix DS-11FX+ Spectrophotometer/ Fluorometer.

### DNA Library Preparation

All PCR reactions were carried out using Kapa HiFi HotStart ready mix 2x Master Mix (KapaBiosystems, cat# 07958935001) in 0.2 mL thin wall PCR plate (Fisher Scientific, AB0800150). The V34 and V4 variable regions were chosen for 16S amplicon sequencing on the fecal samples. V34 (340F (5’→3’) <u>TCGTCGGCAGCGTCAGATGTGTATAAGAGACAG</u>CTACGGGNGGCWGCAG/ 806R (5’→3’) <u>GTCTCGTGGGCTCGGAGATGTGTATAAGAGACAG</u>GGACTACNVGGGTWTCTAAT) and V4 (515F (5’→3’) (<u>TCGTCGGCAGCGTCAGATGTGTATAAGAGACAG</u>GTGYCAGCMGCCGCGGTAA/ 806R (5’→3’) <u>GTCTCGTGGGCTCGGAGATGTGTATAAGAGACAG</u>GGACTACNVGGGTWTCTAAT) were ordered as pre-mixed primer pairs and include the Illumina handles (underlined) on the 5’ end (RxnReady Primer Pools, IDT). The 16S PCR amplification was carried out using the following reaction mixture: 12.5 μL of Kapa HiFi HotStart 2x Master Mix, 1 μL of 10 μM V34 primer pool, 12 ng of extracted DNA, and nuclease free water to bring the final volume to 25 μL. Amplification was carried out using the following protocol: Initial denature (95.0°C – 3:00), 25 cycles of denature (98.0 °C – 0:20), annealing (55.0 °C – 0:15), elongation (72.0 °C – 0:20), and a final extension (72.0 °C – 1:00). The 16S reactions were purified using SPRIselect beads (Beckman Coulter, cat# B23318) to select for amplicons. The beads were added at a 0.8:1 ratio (20 μL of beads for 25 μL reaction), mixed by pipetting, and incubated on the bench for 1 minute then place on magnet for 1 minute. Without disturbing beads, liquid was discarded and while keeping the plate on magnet, 180 µL of 80 % molecular grade ethanol (Sigma-Aldrich, cat# E7023-1L) was added. After 30 seconds, ethanol, was removed carefully to not disturb the beads (first with 200 μL pipette set to 180 μL, then with 20 μL pipette set to 20 μL to remove any excess ethanol. The plate was then removed from magnet and 40 μL of RNase/DNase free molecular biology grade water (Promega, cat# P1195) from was added and mixed by pipetting. Bead were incubated with the water for one minute, the place on magnet and allowed to sit for one minute. Careful not to disturb the beads, water was carefully removed and placed in a new 0.2 mL PCR plate.

Cleaned 16S amplicons were then barcoded using dual indexes from Illumina (Nextera XT Index kit V2, Illumina cat# 15052163) PCR reactions were set up similarly to the 16S reaction described above with the following exceptions, 2.5 μL of cleaned 16S amplicons were added as the DNA template and 5 μL of each indexing primer was used. The same thermocycler conditions were used as for the 16S PCR with the following modification: only 8 cycles were run. Indexed, 16S amplicons were then cleaned following the same SPRIselect protocol detailed above, with the following modifications: beads are added at a 1:1 ratio (25 µL of beads for 25 µL reaction. Following clean-up, samples were quantified by fluorescence (DeNovix dsDNA high sensitivity assay cat# Kit-DSDNA-HIGH-2).

### Sequencing

10 ng of each sample were pooled for the sequencing. The samples were first pooled by operator. Pools were quantified by Fluorescence (DeNovix dsDNA high sensitivity assay cat# Kit-DSDNA-HIGH-2) and nM of each pool was calculated. DNA pools were stored at 4 °C until sequencing. On the day of sequencing, each operator diluted the pools to 4 nM. Pools were combined in equal volume and prepared for sequencing following the MiSeq System Denature and Dilute Libraries Guide (Document # 15039740 v10, Protocol A). Denatured libraries were diluted to a final concentration of 12 pM and combined with 5 % PhiX control (V3 cat# 15017666 from Illumina). Paired-end sequencing was performed on an Illumina MiSeq with 2×300 bp reads (MiSeq Reagent Kit v3 600-cycle, cat #: MS-102-3003).

### Data analysis

Adapter trimming was done as part of the Illumina MiSeq Generate FASTQ workflow. Fastq files were imported to RStudio (R version 4.1.0) for processing. Cutadapt (2.8)(25) was used for primer trimming followed by DADA2 (1.20.0)(26) with taxonomic assignment. Briefly, Cutadapt identified and trimmed primers allowing for no more than 2 mismatches with the primer sequencing. Primer trimmed sequences were then fed into the DADA2 pipeline. Reads were filtered and trimmed using the following parameters: all reads (forwards and reverse) were trimmed to 250 bp, reads could contain no Ns, maxEE (expected error) of five for both forward and reverse reads, and the default trunQ (2). Default settings for error learning, dereplicating, merging, and chimera identification were used. The R code for the initial sequence processing and the statistical analyses shown herein are publicly available at https://data.nist.gov/od/id/mds2-3092.

### NIST Disclaimer

Certain commercial equipment, instruments, or materials are identified in this paper to foster understanding. Such identification does not imply recommendation or endorsement by the National Institute of Standards and Technology, nor does it imply that the materials or equipment identified are necessarily the best available for the purpose. The reference materials used in this study were not certified by NIST and are not official NIST Reference Materials.

### Ethics approval and Consent to Participate

All work was reviewed and approved by the U. S. National Institute of Standards and Technology (NIST) Research Protections Office. This study (protocol #: MML-2019-0135) was determined to be “not human subjects research” as defined in the Common Rule (45 CFR 46, Subpart A).

### Consent for Publication

Not applicable.

### Availability of Data and Materials

All metagenomic sequencing results and the code used for analyses in this manuscript are available online (https://data.nist.gov/od/id/mds2-3092). Remaining units of the fecal materials used for this project are available for purchase from The BioCollective.

### Competing Interests

The authors declare no competing interests.

### Authors Contributions

S.P.F., S.L.S., J.G.K., J.J.F., and S.A.J. conceptualized the project and designed experiments. J.N.D. and M.E.H. analyzed samples. S.P.F. and S.L.S. analyzed data. S.P.F., S.L.S., J.G.K., and S.A.J. wrote the manuscript and prepared figures. All authors approved the final manuscript except for J.J.F. who passed away before the completion of the project.

## Acknowledgements

The authors wish to acknowledge Sheng Lin-Gibson, Kirsten H. Parratt, Alshae R. Logan, and Lisa M. Stabryla for critical feedback during manuscript review.

## Notes

https://data.nist.gov/od/id/mds2-3092

### Competing Interest Statement

The authors have declared no competing interest.

