## Supplemental Figures and Tables for "A Sensitivity Analysis of Methodological Variables Associated with Microbiome Measurements"

**Supporting Information:** to accompany "A Sensitivity Analysis of Methodological Variables Associated with Microbiome Measurements"

**Authors:** Samuel P. Forry, Stephanie L. Servetas, Jennifer N. Dootz, Monique E. Hunter, Jason G. Kralj, James J. Filliben, Scott A. Jackson

**SI Figures and Tables:**

- **SI Figure 1: Stacked taxonomic bar charts to explore methodological choices.** *Alongside Figure 3a from the main manuscript, this taxonomic bar chart is broken out by Sample, Operator, and Extraction Kit. Large differences are observed between the Sample and Extraction Kit parameters (e.g., Coriobacteriaceae), while relatively small differences are observed between Operators (e.g., Akkermansiaceae). However, these notable differences are difficult to quantify. For this plot, Families present at <2% Relative Abundance were grouped as 'Other' (that was 97 taxa across all 5 samples samples).*

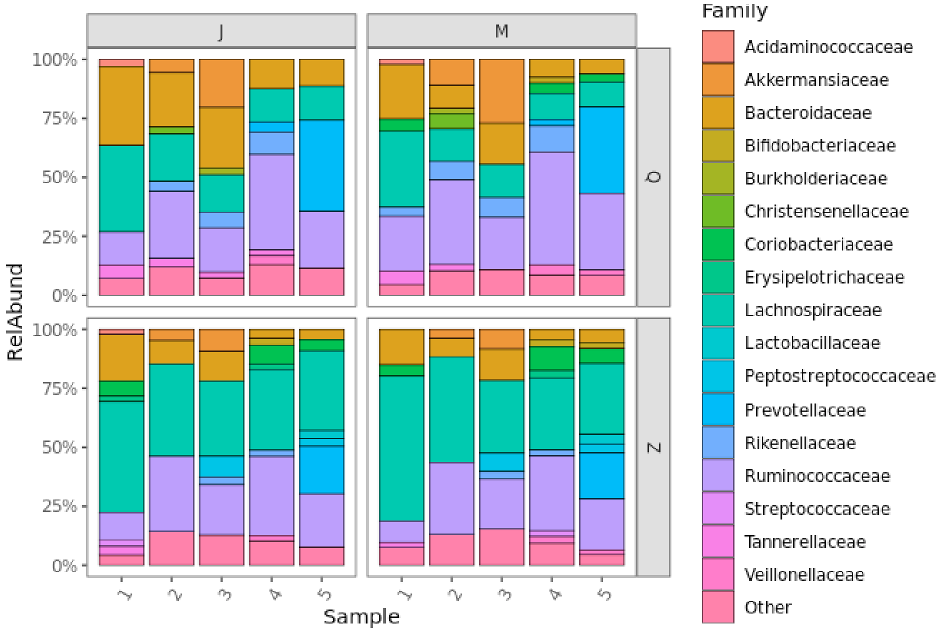

  - 

- **SI Figure 2: Using PCoA plots to explore methodological choices.** *The Bray-Curtis PCoA plot from Figure 3b in the main manuscript was faceted multiple ways to show the impacts of different parameters.* ***SI Figure 2a:*** *The PCoA from Figure 3b of the main manuscript was split by Sample (columns 1-5), while colors and symbols denoted different Extraction Kits.* ***SI Figure 2b:*** *The PCoA plot from Figure SI 2a was split further for each Extraction Kit (rows Q and Z), while colors and symbols denoted different Operators.* ***SI Figure 2c:*** *The PCoA plot from Figure SI 2b was split further for each Operator (sub row J and M), while colors and symbols denoted different material Lots.* ***SI Figure 2d:*** *The PCoA plot from Figure SI 2c was split further Lot (sub-row A and B), while colors and symbols denoted different Variable Regions.* ***SI Figure 2e:*** *The PCoA plot from Figure SI 2d was split further for Variable Region, while colors and symbols denoted different technical Replicates. Only the top and bottom legs of the experimental design (Figure 2, from the main manuscript, orange and blue arrows) were replicated, so other facets of the plot were omitted for clarity.*

○ SI Figure 2a

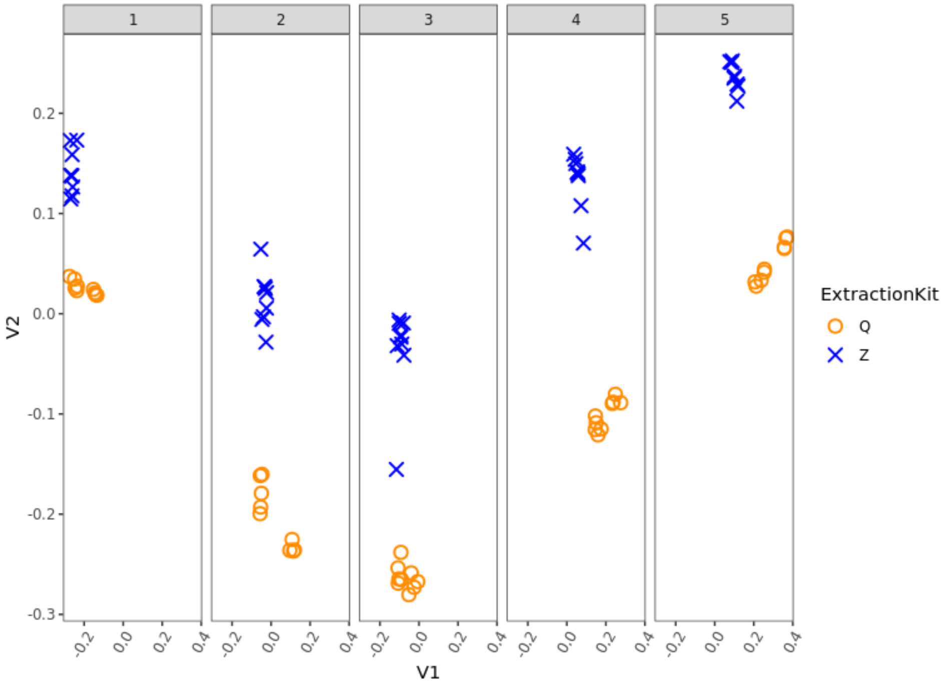

○ SI Figure 2b:

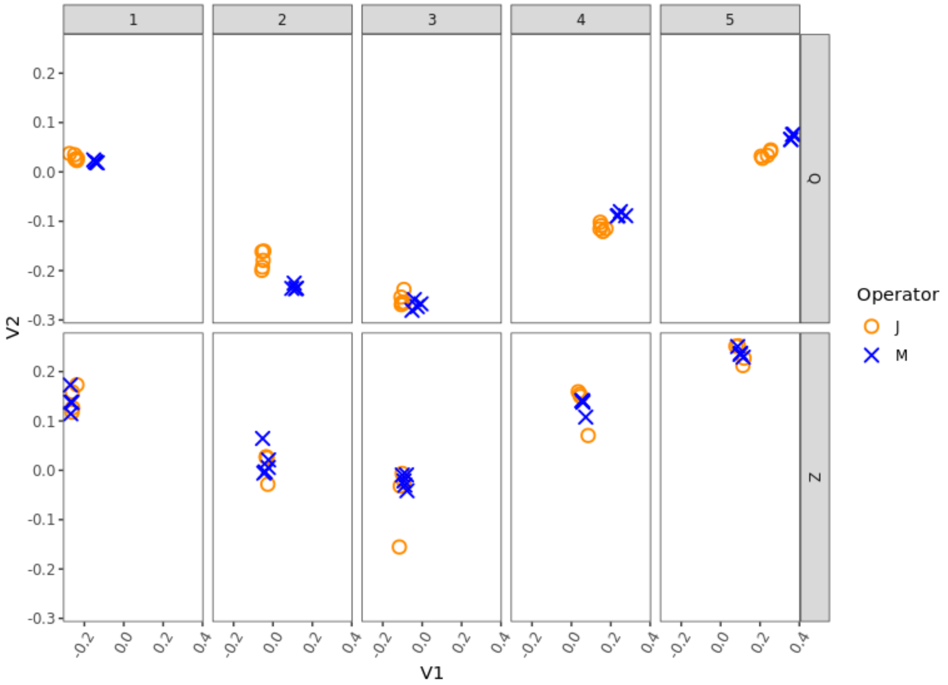

- SI Figure 2c:

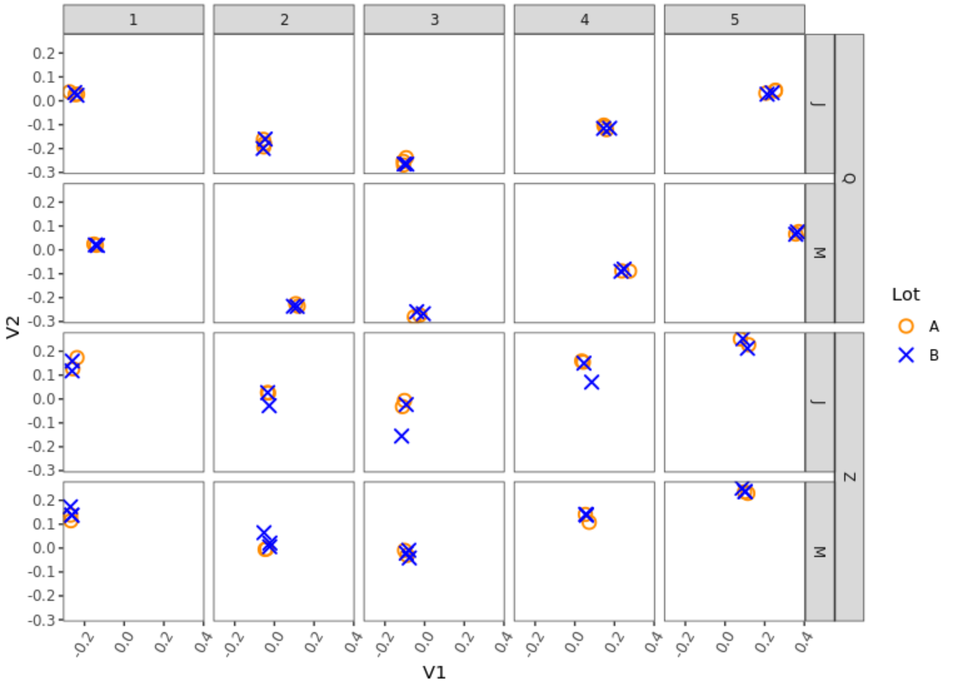

- SI Figure 2d:

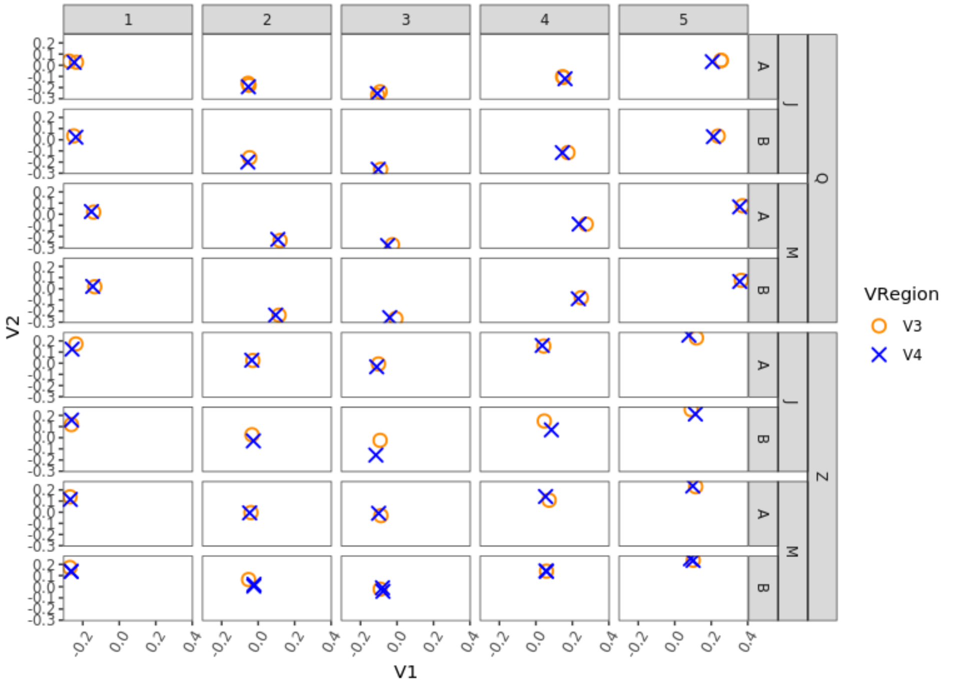

- SI Figure 2e:

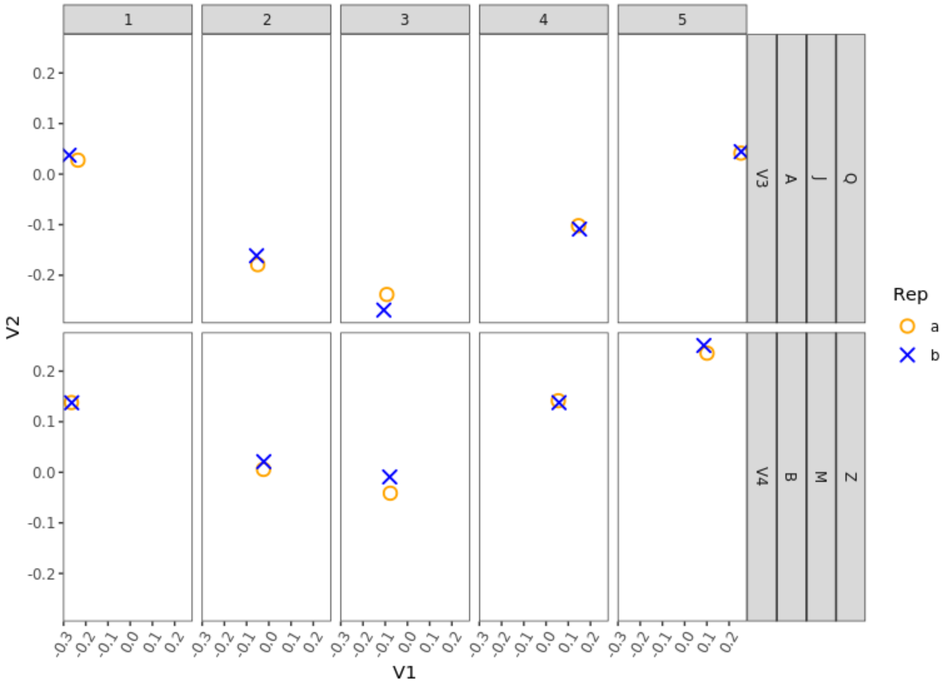

- **Figure SI-3: Protocol choices differentially impact measurements of the two internal standards.** *Parameter Effects were calculated using the ratio Aliivibrio:Leifsonia as described in Equation 1 from the main manuscript, and the Parameter Effects were plotted as a fold change on a log2 scale, such that the horizontal line at 0 denotes the null hypothesis of no effect. As expected, there were no significant biological differences in the ratio of internal standards between samples. However, several methodological choices (e.g., Extraction Kit) did exhibit significant bias between the internal standards. Data error bars showed 99 % confidence intervals, and the points statistically significantly different from the mean (p<0.01) were colored red.*

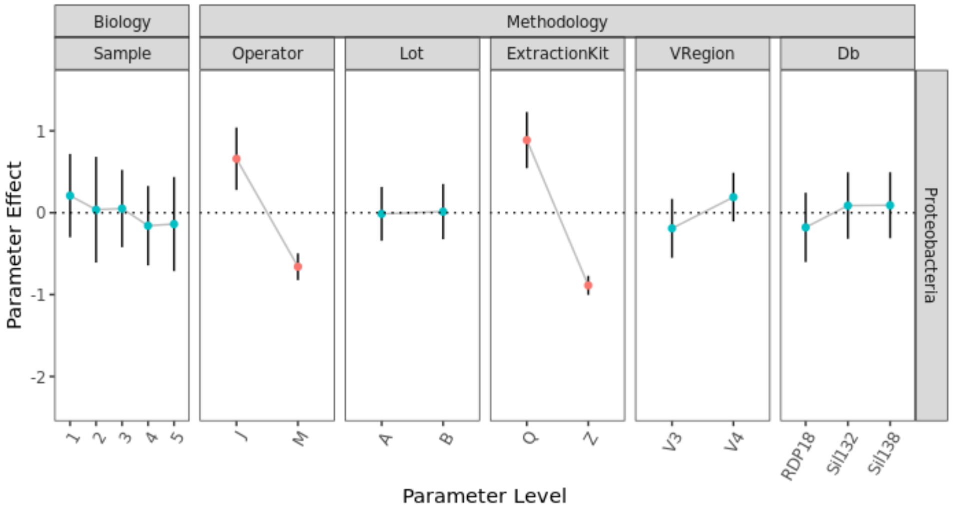

- **Figure SI-4: Parameter Effects for Bacteroidetes within each individual stool sample.** *Alongside Figure 6 from the main manuscript, the Parameter Effects were calculated using the Bacteroidetes:Leifsonia ratio for each individual stool sample (Figure 4a:e for Samples 1:5, respectively). This parameter effect was plotted as a fold change on a log2 scale, such that the*

*horizontal line at 0 denotes the null hypothesis of no effect. The magnitude of the effect of protocol choices (e.g., Extraction Kit) could be directly compared between parameters and parameter levels. Data error bars showed 99 % confidence intervals, and the points that are statistically significantly different from the mean (p<0.01) were colored red.*

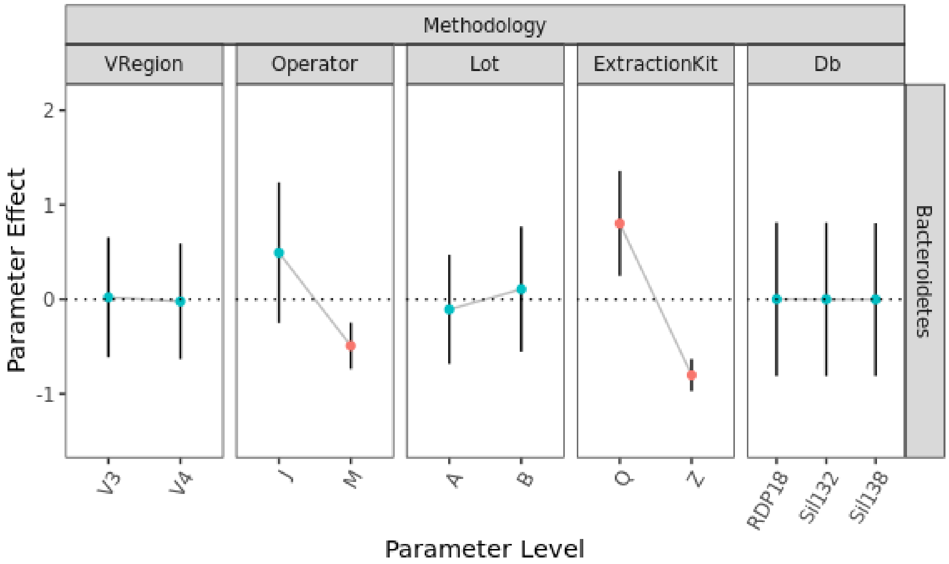

- SI-4A:

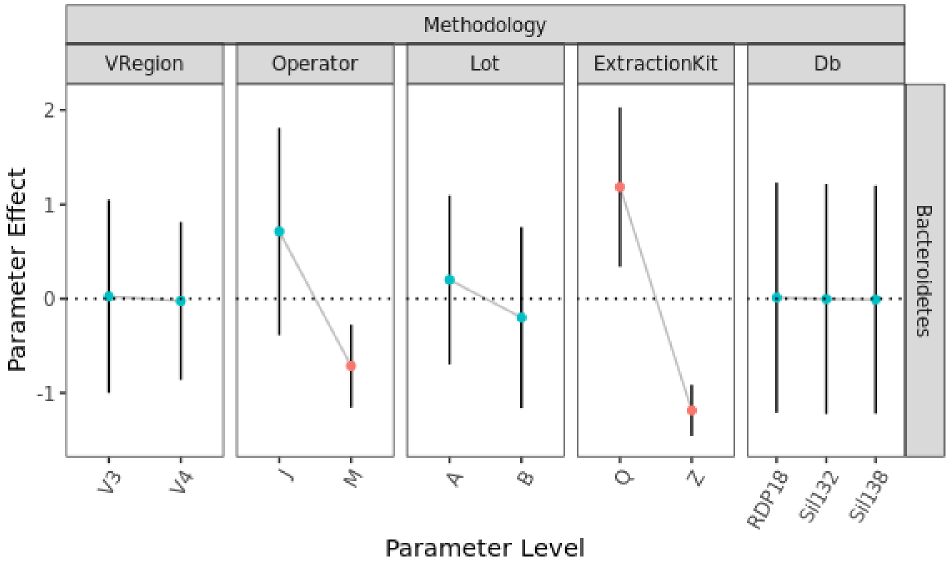

- SI-4B:

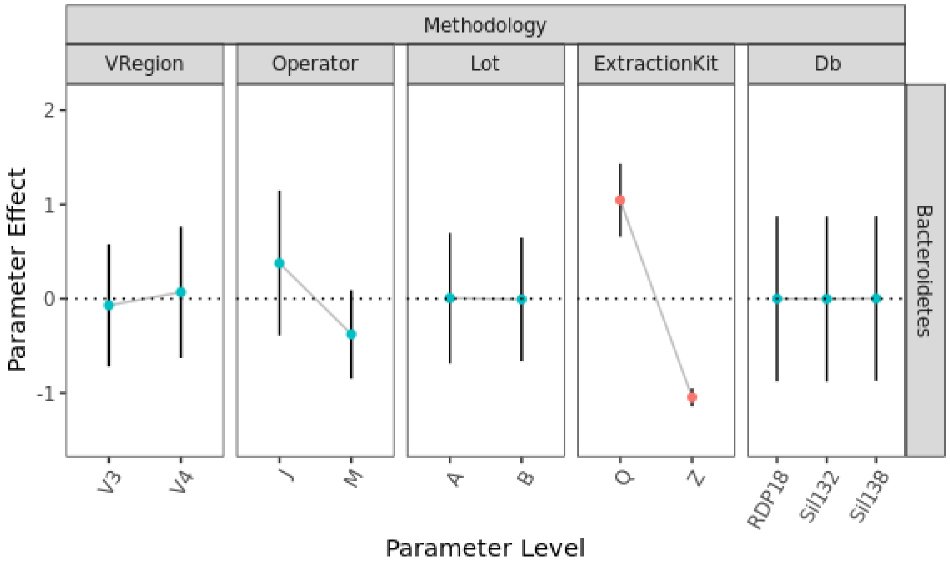
Methodology
VRegion
Operator
Lot
ExtractionKit
Db
Bacteroidetes
Parameter Effect
2
1
0
-1
V3
V4
J
M
A
B
Q
Z
RDP18
Sil132
Sil138
Parameter Level

- SI-4C:

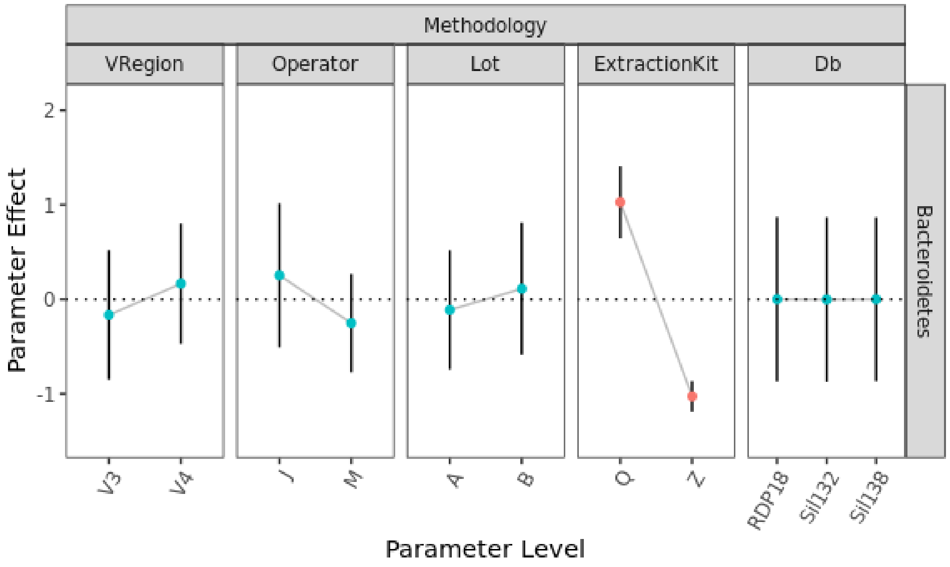
Methodology
VRegion
Operator
Lot
ExtractionKit
Db
Bacteroidetes
Parameter Effect
2
1
0
-1
V3
V4
J
M
A
B
Q
Z
RDP18
Sil132
Sil138
Parameter Level

- SI-4D:

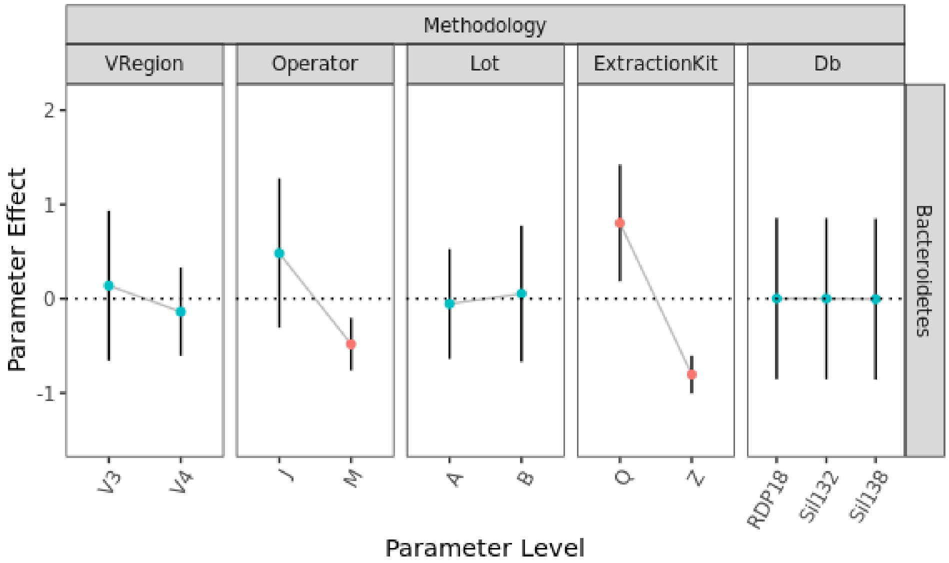

- SI-4E:

- SI Figure SI-5 ***Quantitative comparison of methodological parameters for multiple Phyla*** *. The Parameter Effect was calculated for the ratio of various Phyla of interest to an internal control (Leifsonia) as shown in Equation 1. This parameter effect was plotted as a fold change on a log2 scale, such that the horizontal line at 0 denotes the null hypothesis of no effect. The magnitude of the effect of protocol choices (e.g., Extraction Kit) could be directly compared between parameters and parameter levels. Data error bars showed 99 % confidence intervals, and the points that were statistically significantly different from the mean (p<0.01) were colored red.*

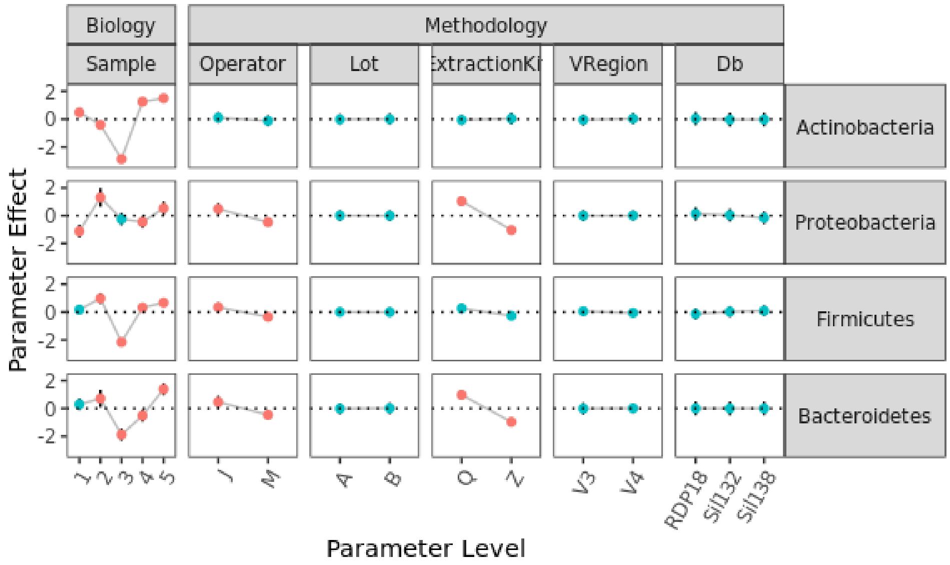

Figure SI-6, ***The effect of methodological parameters varied with taxonomic specificity.*** *The Parameter Effect was calculated for the ratio of various taxa to an internal control (Leifsonia). SI 6a:g correspond to the genera plotted in Figure 7 in the main manuscript. Here the Parameter effects are calculated and plotted for each descending branch of the taxonomic tree (Phlyum, Class, Order, Family, and Genus): SI-6a shows the Bifidobacterium taxonomic lineage; SI-6b shows the Collinsella taxonomic lineage; SI-6c shows the Allistipes taxonomic lineage; SI-6d shows the Anaerostipes taxonomic lineage; SI-6e shows the Blautia taxonomic lineage; SI-6f shows the Faecalibacterium taxonomic lineage; SI-6g shows the Bacteroides taxonomic lineage.*

*Data error bars show a 99 % confidence interval, and statistically significant points (p < 0.01) are colored red.*

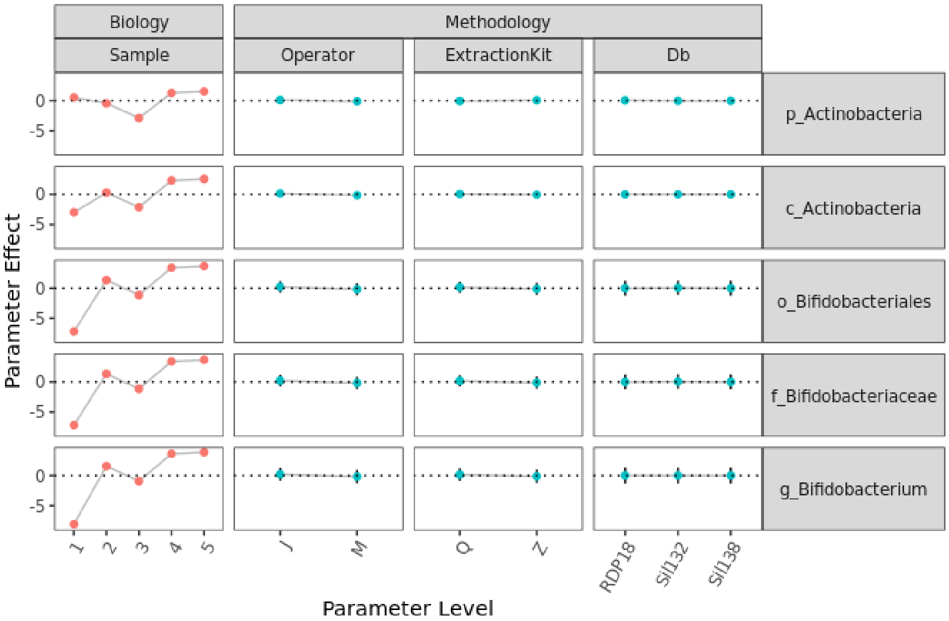

- SI-6a:

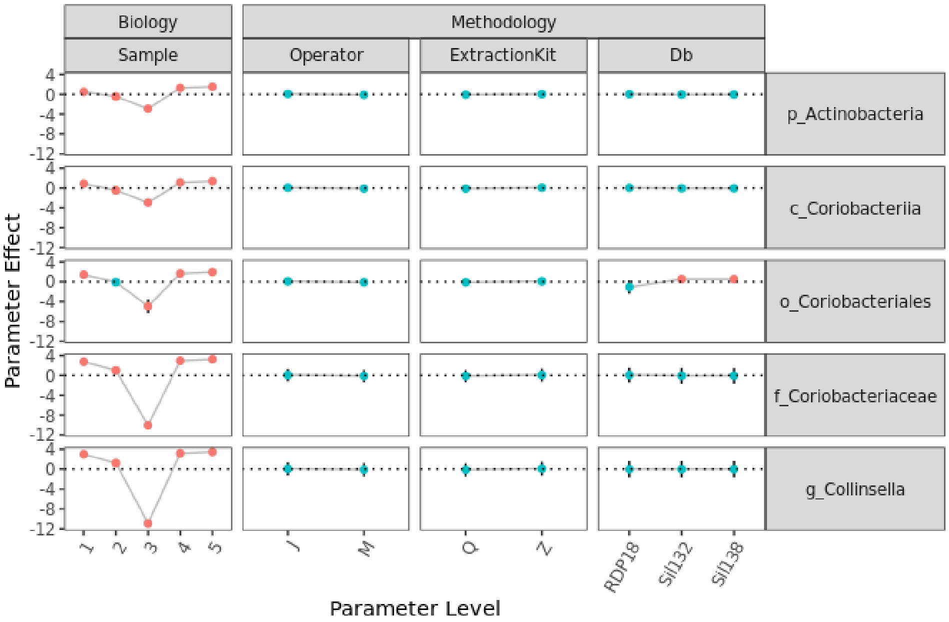

- SI 6b:

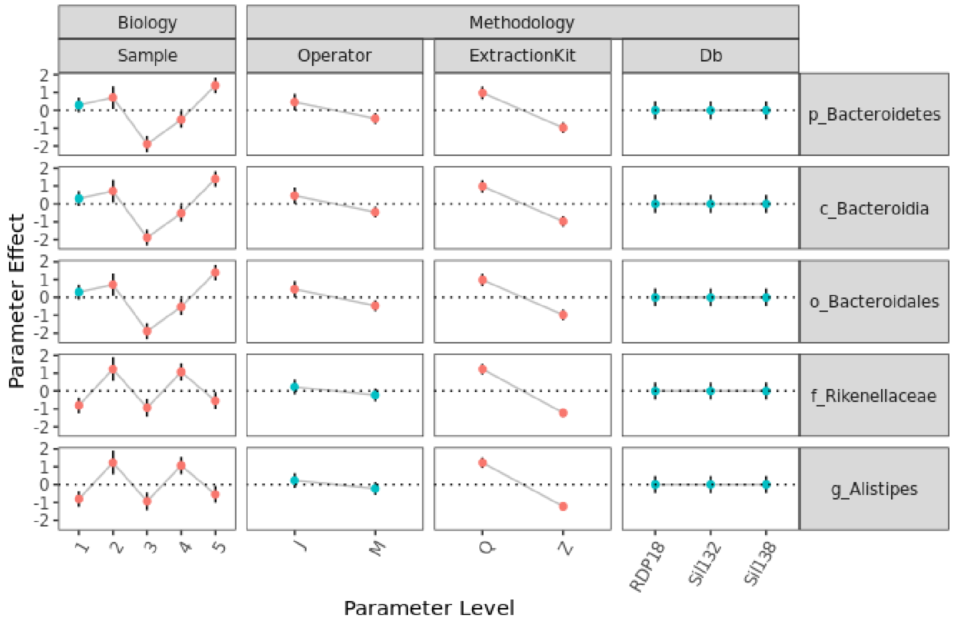
Biology
Methodology
Sample
Operator
ExtractionKit
Db
p_Bacteroidetes
c_Bacteroidia
o_Bacteroidales
f_Rikenellaceae
g_Alistipes
Parameter Effect
2
1
0
-1
-2
1
2
3
4
5
J
M
Q
Z
RDP18
Sil132
Sil138
Parameter Level

- SI 6c:

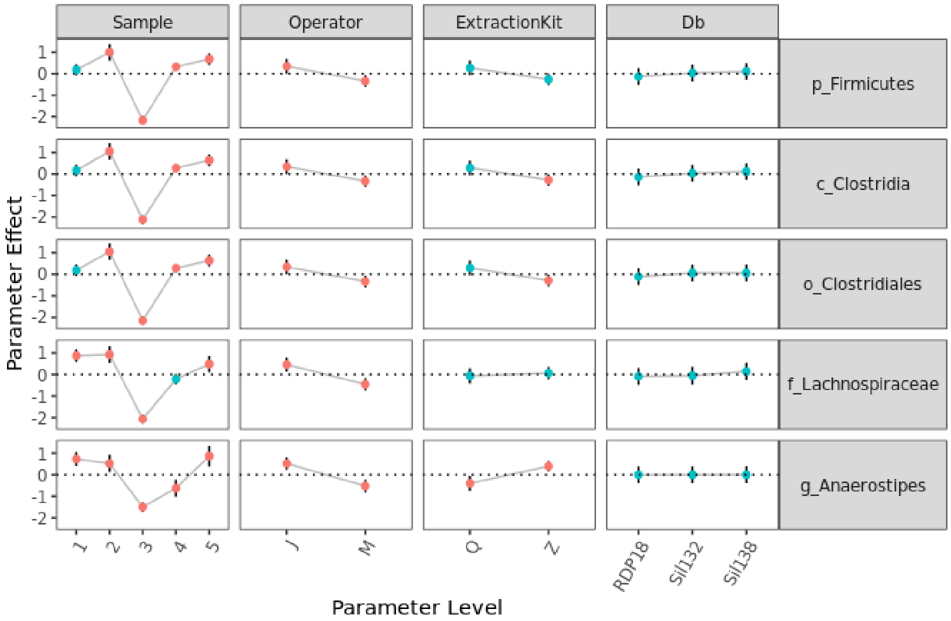
Sample
Operator
ExtractionKit
Db
p_Firmicutes
c_Clostridia
o_Clostridiales
f_Lachnospiraceae
g_Anaerostipes
Parameter Effect
1
0
-1
-2
1
2
3
4
5
J
M
Q
Z
RDP18
Sil132
Sil138
Parameter Level

- SI 6d:

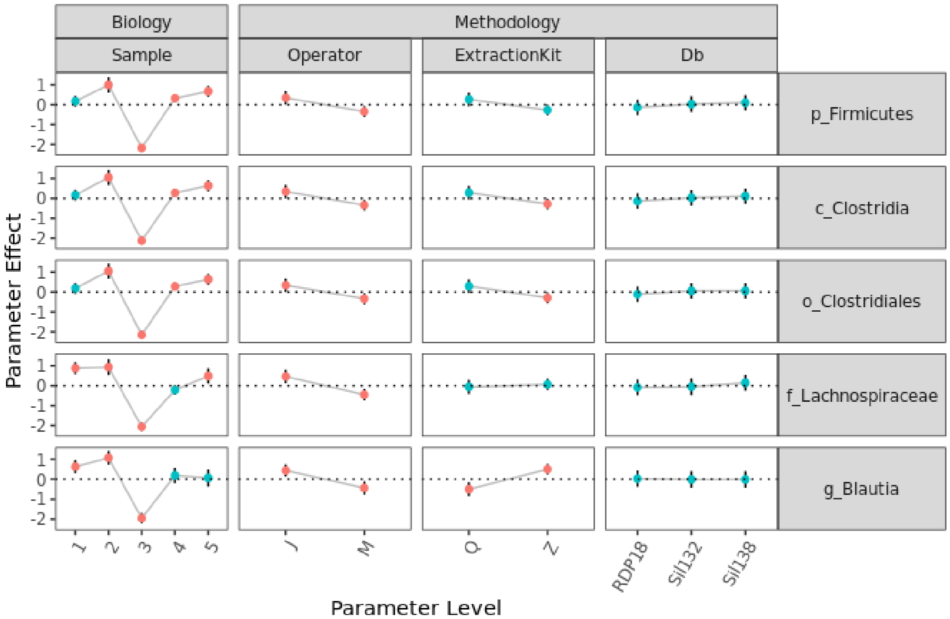
Biology
Methodology
Sample
Operator
ExtractionKit
Db
p_Firmicutes
c_Clostridia
o_Clostridiales
f_Lachnospiraceae
g_Blautia
Parameter Effect
Parameter Level
RDP18
Sil132
Sil138

- SI 6e:

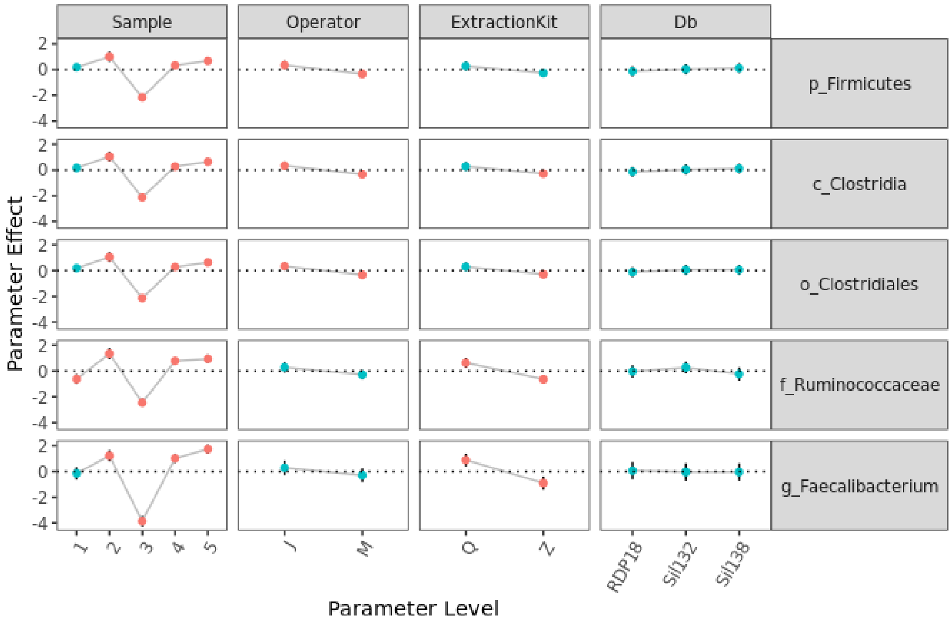
Sample
Operator
ExtractionKit
Db
p_Firmicutes
c_Clostridia
o_Clostridiales
f_Ruminococcaceae
g_Faecalibacterium
Parameter Effect
Parameter Level
RDP18
Sil132
Sil138

- SI 6f:

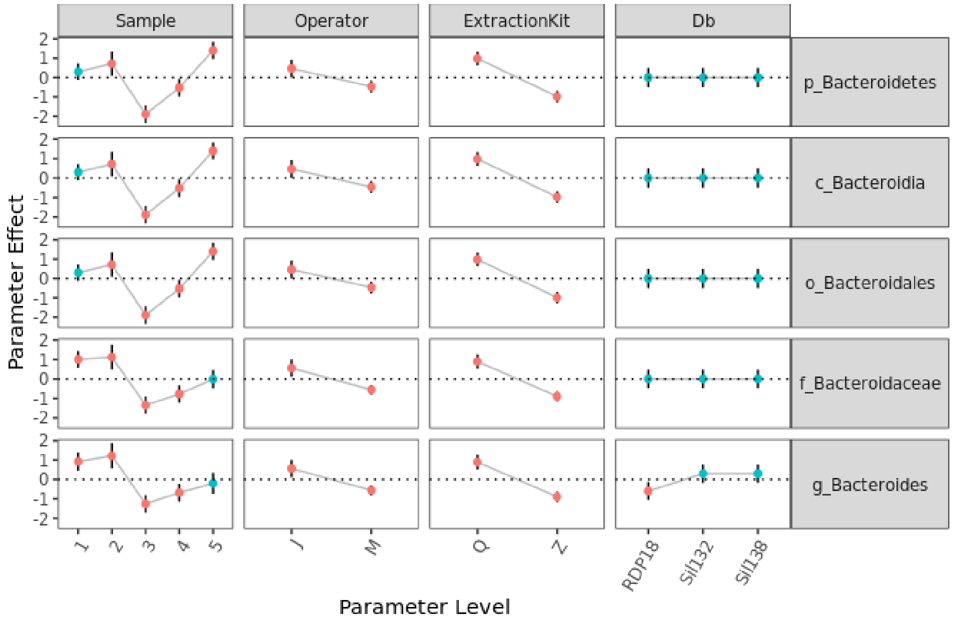

- SI 6g:

- **SI Table 1. Taxonomy Key.** *The complete taxonomy (phyla, class, order, family) of each genus plotted in Figure 8 of the main manuscript is provided here.*

| P | Phylum | C | Class | O | Order | F | Family |
|---|---|---|---|---|---|---|---|
| 1 | Actinobacteria | 1 | Actinobacteria | 1 | Actinomycetales | 1 | Actinomycetaceae |
| 1 | Actinobacteria | 1 | Actinobacteria | 2 | Bifidobacteriales | 2 | Bifidobacteriaceae |
| 1 | Actinobacteria | 1 | Actinobacteria | 3 | Corynebacteriales | 3 | Corynebacteriaceae |
| 1 | Actinobacteria | 1 | Actinobacteria | 4 | Micrococcales | 4 | Microbacteriaceae |
| 1 | Actinobacteria | 1 | Actinobacteria | 4 | Micrococcales | 5 | Micrococcaceae |
| 1 | Actinobacteria | 1 | Actinobacteria | 5 | Propionibacteriales | 6 | Nocardioidaceae |
| 1 | Actinobacteria | 1 | Actinobacteria | 5 | Propionibacteriales | 7 | Propionibacteriaceae |
| 1 | Actinobacteria | 2 | Coriobacteriia | 6 | Coriobacteriales | 8 | Atopobiaceae |
| 1 | Actinobacteria | 2 | Coriobacteriia | 6 | Coriobacteriales | 9 | Coriobacteriaceae |
| 1 | Actinobacteria | 2 | Coriobacteriia | 6 | Coriobacteriales | 10 | Coriobacteriales Incertae Sedis |
| 1 | Actinobacteria | 2 | Coriobacteriia | 6 | Coriobacteriales | 11 | Eggerthellaceae |
| 2 | Bacteroidetes | 3 | Bacteroidia | 7 | Bacteroidales | 12 | Bacteroidaceae |
| 2 | Bacteroidetes | 3 | Bacteroidia | 7 | Bacteroidales | 13 | Barnesiellaceae |
| 2 | Bacteroidetes | 3 | Bacteroidia | 7 | Bacteroidales | 14 | Marinifilaceae |
| 2 | Bacteroidetes | 3 | Bacteroidia | 7 | Bacteroidales | 15 | Porphyromonadaceae |
| 2 | Bacteroidetes | 3 | Bacteroidia | 7 | Bacteroidales | 16 | Prevotellaceae |
| 2 | Bacteroidetes | 3 | Bacteroidia | 7 | Bacteroidales | 17 | Rikenellaceae |
| 2 | Bacteroidetes | 3 | Bacteroidia | 7 | Bacteroidales | 18 | Tannerellaceae |
| 2 | Bacteroidetes | 3 | Bacteroidia | 8 | Flavobacteriales | 19 | Flavobacteriaceae |
| 3 | Campilobacterota | 4 | Campylobacteria | 9 | Campylobacterales | 20 | Campylobacteraceae |
| 4 | Cyanobacteria | 5 | Cyanobacteriia | 10 | Cyanobacteriales | 21 | Phormidiaceae |
| 5 | Deinococcota | 6 | Deinococci | 11 | Deinococcales | 22 | Deinococcaceae |

| | | | | | | | |
|---|---|---|---|---|---|---|---|
| 6 | Desulfobacterota | 7 | Desulfovibrionia | 12 | Desulfovibrionales | 23 | Desulfovibrionaceae |
| 7 | Euryarchaeota | 8 | Methanobacteria | 13 | Methanobacteriales | 24 | Methanobacteriaceae |
| 8 | Firmicutes | 9 | Bacilli | 14 | Bacillales | 25 | Bacillaceae |
| 8 | Firmicutes | 9 | Bacilli | 14 | Bacillales | 26 | Planococcaceae |
| 8 | Firmicutes | 9 | Bacilli | 15 | Erysipelotrichales | 27 | Erysipelatoclostridiaceae |
| 8 | Firmicutes | 9 | Bacilli | 15 | Erysipelotrichales | 28 | Erysipelotrichaceae |
| 8 | Firmicutes | 9 | Bacilli | 16 | Lactobacillales | 29 | Aerococcaceae |
| 8 | Firmicutes | 9 | Bacilli | 16 | Lactobacillales | 30 | Carnobacteriaceae |
| 8 | Firmicutes | 9 | Bacilli | 16 | Lactobacillales | 31 | Enterococcaceae |
| 8 | Firmicutes | 9 | Bacilli | 16 | Lactobacillales | 32 | Lactobacillaceae |
| 8 | Firmicutes | 9 | Bacilli | 16 | Lactobacillales | 33 | Leuconostocaceae |
| 8 | Firmicutes | 9 | Bacilli | 16 | Lactobacillales | 34 | Streptococcaceae |
| 8 | Firmicutes | 9 | Bacilli | 17 | Staphylococcales | 35 | Gemellaceae |
| 8 | Firmicutes | 9 | Bacilli | 17 | Staphylococcales | 36 | Staphylococcaceae |
| 8 | Firmicutes | 10 | Clostridia | 18 | Christensenellales | 37 | Christensenellaceae |
| 8 | Firmicutes | 10 | Clostridia | 19 | Clostridiales | 38 | Clostridiaceae |
| 8 | Firmicutes | 10 | Clostridia | 20 | Eubacteriales | 39 | Anaerofustaceae |
| 8 | Firmicutes | 10 | Clostridia | 20 | Eubacteriales | 40 | Eubacteriaceae |
| 8 | Firmicutes | 10 | Clostridia | 21 | Lachnospirales | 41 | Defluviitaleaceae |
| 8 | Firmicutes | 10 | Clostridia | 21 | Lachnospirales | 42 | Lachnospiraceae |
| 8 | Firmicutes | 10 | Clostridia | 22 | Monoglobales | 43 | Monoglobaceae |
| 8 | Firmicutes | 10 | Clostridia | 23 | Oscillospirales | 44 | Butyricicoccaceae |
| 8 | Firmicutes | 10 | Clostridia | 23 | Oscillospirales | 45 | Ethanoligenenaceae |
| 8 | Firmicutes | 10 | Clostridia | 23 | Oscillospirales | 46 | Oscillospiraceae |
| 8 | Firmicutes | 10 | Clostridia | 23 | Oscillospirales | 47 | Ruminococcaceae |
| 8 | Firmicutes | 10 | Clostridia | 24 | Peptococcales | 48 | Peptococcaceae |
| 8 | Firmicutes | 10 | Clostridia | 25 | Peptostreptococcales-Tissierellales | 49 | Anaerovoracaceae |
| 8 | Firmicutes | 10 | Clostridia | 25 | Peptostreptococcales-Tissierellales | 50 | Peptostreptococcaceae |
| 8 | Firmicutes | 11 | Negativicutes | 26 | Acidaminococcales | 51 | Acidaminococcaceae |
| 8 | Firmicutes | 11 | Negativicutes | 27 | Veillonellales-Selenomonadales | 52 | Selenomonadaceae |
| 8 | Firmicutes | 11 | Negativicutes | 27 | Veillonellales-Selenomonadales | 53 | Sporomusaceae |
| 8 | Firmicutes | 11 | Negativicutes | 27 | Veillonellales-Selenomonadales | 54 | Veillonellaceae |
| 8 | Firmicutes | 12 | Syntrophomonadia | 28 | Syntrophomonadales | 55 | Syntrophomonadaceae |
| 9 | Fusobacteriota | 13 | Fusobacteriia | 29 | Fusobacteriales | 56 | Fusobacteriaceae |
| 10 | Patescibacteria | 14 | Saccharimonadia | 30 | Saccharimonadales | 57 | Saccharimonadaceae |
| 11 | Proteobacteria | 15 | Alphaproteobacteria | 31 | Caulobacterales | 58 | Caulobacteraceae |
| 11 | Proteobacteria | 15 | Alphaproteobacteria | 32 | Rhizobiales | 59 | Beijerinckiaceae |
| 11 | Proteobacteria | 15 | Alphaproteobacteria | 33 | Rhodobacterales | 60 | Rhodobacteraceae |
| 11 | Proteobacteria | 15 | Alphaproteobacteria | 34 | Sphingomonadales | 61 | Sphingomonadaceae |
| 11 | Proteobacteria | 16 | Gammaproteobacteria | 35 | Aeromonadales | 62 | Aeromonadaceae |

| | | | | | | | |
|---|---|---|---|---|---|---|---|
| 11 | Proteobacteria | 16 | Gammaproteobacteria | 36 | Alteromonadales | 63 | Alteromonadaceae |
| 11 | Proteobacteria | 16 | Gammaproteobacteria | 37 | Burkholderiales | 64 | Alcaligenaceae |
| 11 | Proteobacteria | 16 | Gammaproteobacteria | 37 | Burkholderiales | 65 | Burkholderiaceae |
| 11 | Proteobacteria | 16 | Gammaproteobacteria | 37 | Burkholderiales | 66 | Comamonadaceae |
| 11 | Proteobacteria | 16 | Gammaproteobacteria | 37 | Burkholderiales | 67 | Neisseriaceae |
| 11 | Proteobacteria | 16 | Gammaproteobacteria | 37 | Burkholderiales | 68 | Oxalobacteraceae |
| 11 | Proteobacteria | 16 | Gammaproteobacteria | 37 | Burkholderiales | 69 | Rhodocyclaceae |
| 11 | Proteobacteria | 16 | Gammaproteobacteria | 37 | Burkholderiales | 70 | Sutterellaceae |
| 11 | Proteobacteria | 16 | Gammaproteobacteria | 38 | Cardiobacteriales | 71 | Cardiobacteriaceae |
| 11 | Proteobacteria | 16 | Gammaproteobacteria | 39 | Cellvibrionales | 72 | Cellvibrionaceae |
| 11 | Proteobacteria | 16 | Gammaproteobacteria | 40 | Enterobacterales | 73 | Enterobacteriaceae |
| 11 | Proteobacteria | 16 | Gammaproteobacteria | 40 | Enterobacterales | 74 | Yersiniaceae |
| 11 | Proteobacteria | 16 | Gammaproteobacteria | 41 | Pasteurellales | 75 | Pasteurellaceae |
| 11 | Proteobacteria | 16 | Gammaproteobacteria | 42 | Pseudomonadales | 76 | Moraxellaceae |
| 11 | Proteobacteria | 16 | Gammaproteobacteria | 42 | Pseudomonadales | 77 | Pseudomonadaceae |
| 11 | Proteobacteria | 16 | Gammaproteobacteria | 43 | Vibrionales | 78 | Vibrionaceae |
| 11 | Proteobacteria | 16 | Gammaproteobacteria | 44 | Xanthomonadales | 79 | Xanthomonadaceae |
| 12 | Synergistota | 17 | Synergistia | 45 | Synergistales | 80 | Synergistaceae |
| 13 | Thermoplasmatota | 18 | Thermoplasmata | 46 | Methanomassiliicoccales | 81 | Methanomassiliicoccaceae |
| 14 | Verrucomicrobia | 19 | Lentisphaeria | 47 | Victivallales | 82 | Victivallaceae |
| 14 | Verrucomicrobia | 20 | Verrucomicrobiae | 48 | Verrucomicrobiales | 83 | Akkermansiaceae |
